# Mitotic tethering enables *en masse* inheritance of a shattered micronuclear chromosome

**DOI:** 10.1101/2022.09.08.507171

**Authors:** Prasad Trivedi, Christopher D. Steele, Ludmil B. Alexandrov, Don W. Cleveland

## Abstract

Chromothripsis, the shattering and imperfect reassembly of one (or a few) chromosome(s)^1^, is an ubiquitous^2^ mutational process generating localized complex chromosomal rearrangements that drive genome evolution in cancer. Chromothripsis can be initiated by missegregation errors in mitosis^3,4^ or DNA metabolism^5-7^ that lead to entrapment of chromosomes within micronuclei and their subsequent fragmentation in the next interphase or upon mitotic entry^6,8-10^. Here, we use inducible degrons to demonstrate that chromothriptically produced pieces of a micronucleated chromosome are tethered together in mitosis by a protein complex consisting of Mediator of DNA damage checkpoint 1 (MDC1), DNA Topoisomerase II Binding Protein 1 (TOPBP1), and Cellular Inhibitor of PP2A (CIP2A), thereby enabling *en masse* segregation to the same daughter cell. Such tethering is shown to be crucial for the viability of cells undergoing chromosome missegregation and shattering after transient inactivation of the spindle assembly checkpoint. Pan-cancer analysis of transcriptome and whole-genome sequencing data revealed that chromothriptic genome rearrangements were accompanied by elevated expression of MDC1, TOPBP1, and CIP2A. Thus, chromatin-bound tethers maintain proximity of fragments of a shattered chromosome enabling their re-encapsulation into, and religation within, a daughter cell nucleus to form heritable, chromothriptically rearranged chromosomes found in the majority of human cancers.

## Introduction

Elevated rates of chromosome missegregation, caused by errors in mitosis, are frequently observed in many cancers^11^. Chromosome missegregation can lead to numerical and structural chromosomal changes in cancer cells, resulting in increased phenotypic diversity, which provides a substrate for natural selection^11^. Defects in chromosome segregation can result in abnormal structures including micronuclei or chromatin bridges, which are prevalent across the spectrum of human neoplasia. Micronuclei and chromatin bridge formation trigger chromothripsis^1,3-8,12^, an event in which a chromosome is shattered into ten to hundreds of individual pieces, most of which are acentric. This shattering is followed by re-ligation in random order of individual pieces, with a few pieces missing, resulting in a massively rearranged chromosome(s). Such chromothriptic rearrangements can result in multiple oncogenic insults, including loss of tumor suppressors^13^, amplification of an oncogenes^1,13-15^, and formation of oncogenic fusions^15^. Chromothripsis can also drive formation and evolution of circular extrachromosomal DNA (ecDNA)^12^, a major vehicle for oncogene amplification. How the pieces of a shattered chromosome are brought together and ligated to yield a chromothriptic chromosome is unknown^16^. Nevertheless, following micronucleation, newly generated structural variants are preferentially present in one of the two daughter cells^3^. Here, we identify a complex of MDC1, TOPBP1, and CIP2A whose recruitment to *γ*H2AX-containing chromatin is responsible for tethering shattered chromosomal fragments to each other, thereby enabling their delivery *en masse* to a daughter cell nucleus and reassembly into a chromothriptically rearranged chromosome.

## Results

### The fragments of a shattered chromosome are clustered in mitosis

To understand the behavior of a broken chromosome resulting from encapsulation into a micronucleus or a chromatin bridge, we generated fragmented chromosomes in RPE1 p53^-/-^ cells using two approaches. First, we added an inhibitor (NMS-P715) of the mitotic checkpoint kinase Mps1 to induce chromosome missegregation and micronuclei formation^17^. Second, low dose inhibition of Topoisomerase II (with ICRF-193) was used to generate catenated sister chromatids to produce mitotic chromatin bridges^7^, which would subsequently fragment at or after cytokinesis (**Fig. 1*A***). Since chromosomes within micronuclei undergo fragmentation before or upon entry into mitosis^6-9^, cells were arrested in the next mitosis using an inhibitor of the mitotic kinesin Eg5 to chronically activate the spindle assembly checkpoint. To follow the fragmented chromosome pieces, we exploited that an initial DNA damage response to a double stranded break is the replacement of histone H2A with the *γ*H2AX variant throughout ∼1 Mb of chromatin^18-20^ adjacent to the break. Generation of chromosome fragmentation by either method yielded *γ*H2AX-containing chromosome fragments that remained tightly clustered in mitosis for large majorities (75.5%±0.5 or 65.5%±0.7, respectively) of fragmented chromosomes arising from Mps1 or Topoisomerase inhibition (**Fig. 1*B, C***).

**Fig. 1.**
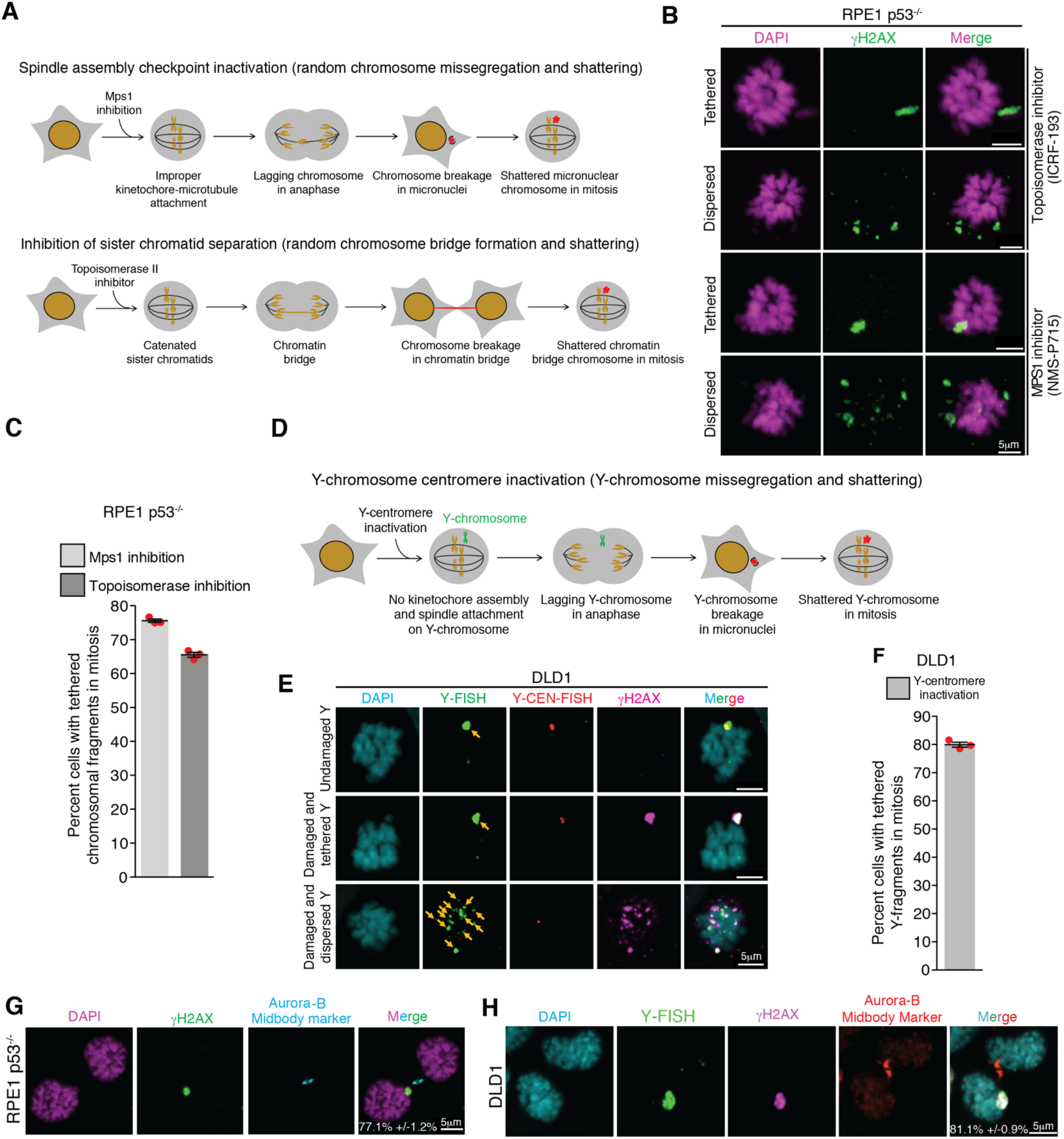
Chromosome fragments from abnormal nuclear structures are clustered during mitosis and segregated together to a daughter cell. ***(A)*** Schematic of methods for generation of micronuclei and chromatin bridges in RPE1 p53^-/-^ cells. ***(B)*** Representative images showing dispersed and tethered chromosomal fragments after micronuclei or chromatin bridge induction. ***(C)*** Quantitation of chromosomal fragment tethering from experiments in *(A)* and *(B)* (n=3 independent experiments, total 511 and 420 cells were analyzed for micronuclei and chromatin bridge induced condition, respectively). ***(D)*** Schematic for generation of Y-chromosome micronuclei in DLD1 cells. ***(E)*** Representative images showing different fates of micronuclear Y-chromosome in mitosis (yellow arrows point to the Y-chromosome fragments) (Y-FISH is Y-chromosome FISH and Y-CEN-FISH is Y-centromere FISH). ***(F)*** Quantitation of tethered Y-chromosome fragments during mitosis 3 days post Y-centromere inactivation (n=3 independent experiments, total 252 cells were analyzed). Representative images showing asymmetric inheritance of broken micronuclear chromosomes in ***(G)*** RPE1 p53^-/-^ cells (n=3 independent experiments, total 201 were analyzed) and ***(H)*** DLD1 cells (n=3 independent experiments, total 143 were analyzed) (for (*G)* and (*H)*, percentage of cells (mean +/- standard error) showing *en masse* inheritance of chromosomal fragments is indicated on the image). For graphs shown in *(C)* and *(F)*, the mean and standard error are shown; individual data points are shown as solid red circles.

Next, we utilized a DLD1 cell model^8^ in which missegregation and micronucleation of the Y-chromosome could be triggered by selective inactivation of its centromere (**Fig. 1*D***). As expected, Y-centromere inactivation resulted in damaged, *γ*H2AX positive Y-chromosomes in the next mitosis, as determined by combined immunofluorescence for *γ*H2AX and fluorescence *in situ* hybridization (FISH) for the Y-chromosome (**Fig. 1*E***) or by Y-chromosome FISH on chromosome spreads (**Extended Data Fig. 1*A-C***). The proportion of cells with a *γ*H2AX positive Y-chromosome (**Extended Data Fig. 2*D***) was similar to the proportion of fragmented Y-chromosomes (**Extended Data Fig. 1*C***), confirming *γ*H2AX to be a reliable marker for fragmented mitotic chromosomes. Once again, in almost all (79%±0.9) cells,*γ*H2AX positive, Y-chromosome fragments remain clustered even in extended mitoses (**Fig. 1*E, F***). Asymmetric segregation of clustered fragments from an initially micronucleated chromosome to one daughter cell in the subsequent G1 phase was consistently observed, both in RPE1 p53^-/-^ cells and in DLD1 cells (**Fig. 1*G, H***). Thus, shattered chromosome fragments produced from micronuclei or chromatin bridges remained clustered in the subsequent mitosis.

### Fragment clustering in mitosis is not due to persistence of the micronuclear envelope

Nuclear envelope disassembly of some micronuclei can be delayed compared to the primary nucleus^4^. Correspondingly, we examined the nuclear envelope around clustered micronuclear chromosome fragments in mitosis after induction of Y-chromosome micronucleation. No damaged Y-chromosome had detectable nuclear envelope remaining (assessed by lamin A/C or lamin B1 staining - **Extended Data Fig. 2*A, B***). This result eliminates the persistence of the micronuclear envelope as an explanation for the clustering of micronuclear chromosomal fragments in mitosis.

### PolD3, MRN complex, and PolQ do not mediate fragment clustering

DNA replication in micronuclei is frequently delayed compared to the primary nucleus, yielding micronuclear DNA that is incompletely replicated at mitotic entry^9,10^. Under-replicated regions are known to undergo PolD3- (a subunit of polymerase Delta) mediated mitotic DNA repair synthesis (MIDAS)^21^. Chromosomes from micronuclei or an initial chromatin bridge have been reported to undergo a burst of DNA synthesis during mitosis^7^. We tested if replication intermediates formed as a result of PolD3-mediated mitotic DNA synthesis could facilitate the clustering of chromosomal pieces in mitosis. The percentage of cells with a damaged Y-chromosome was measured in the subsequent mitosis after depletion of PolD3 with siRNA and the triggering of Y-chromosome micronucleation (**Extended Data Fig. 2*C, D***). Although depletion of PolD3 or the key non-homologous end joining (NHEJ) ligase, Lig4, doubled the percentage of cells with a broken mitotic Y-chromosome (**Extended Data Fig. 2*E***), it did not alter the proportion of cells with clustered micronuclear chromosomal fragments (**Extended Data Fig. 2*F***). PolD3 involvement in damaged micronuclear chromosome repair was confirmed (**Extended Data Fig. 2*G-J***) using “CEN-select”, a colony formation assay to monitor reassembly of a micronuclear chromosome by selecting for retention of a neomycin resistance gene inserted into an initially intact Y-chromosome^4^. Thus, PolD3 contributes to repair of fragmented micronuclear chromosomes, but does not play a role in mitotic clustering of the micronuclear chromosomal fragments.

The MRX (MRN in humans) complex consisting of Mre11, Rad50, and Xrs2 (Nbs1 in humans) has been reported to play a role in tethering an acentric piece of a broken chromosome to the centromere-containing fragment during mitosis in yeast^22,23^ and flies^24^. Indeed, human MRN can tether DNA ends *in vitro*^*25*^. We tested if the MRN complex mediates tethering of micronuclear fragments during mitosis by inducing Y-chromosome micronucleation after prior depletion of either of two members (Mre11 or Nbs1) of the MRN complex using siRNA (**Extended Data Fig. 3*A***). MRN complex depletion did not affect the frequency of cells with tethered Y-pieces (**Extended Data Fig. 3*B-D***). Similarly, depletion of PolQ, reported to promote segregation of an acentric chromosome fragment in Drosophila^24^, did not alter Y-chromosomal fragment tethering (**Extended Data Fig. 3*E***). Thus, mitotic clustering of fragments from a micronucleated chromosome is not mediated by the MRN complex or PolQ.

### MDC1, TOPBP1, and CIP2A localize to broken micronuclear chromosomes at mitotic entry

Similar to the MRN complex and PolQ, TOPBP1 has also been proposed to maintain chromosomal integrity in mitosis after DNA damage resulting from ionizing radiation (IR)^26,27^ or replication stress caused by BRCA1/2 deficiency^28^. TOPBP1 is known to be recruited to some DNA damage lesions in mitosis through direct interaction with MDC1^27^, which in turn directly binds to the mega-base domains of *γ*H2AX-containing chromatin assembled at DSBs^18^. CIP2A interacts with TOPBP1 and mediates its recruitment to DNA damage sites in mitosis^26,28,29^. Both ruptured and intact micronuclei have abnormal nucleoplasm (in the latter case as a consequence of deficient nuclear import^9,30^) and during interphase they fail to recruit key DNA repair proteins, including 53BP1 and BRCA1, even after DNA damage marked by *γ*H2AX (**Extended Data Fig. 3*F, G***). While we determined that a low level of MDC1, TOPBP1, or CIP2A was accumulated within micronuclei containing damaged micronuclear chromosomes in a minority (23.7±2.0%, 12.0±1.4%, and 17.6±0.3%, respectively) of interphase DLD1 cells and a similar number for RPE1 p53^-/-^ cells (**Extended Data Fig. 4**A-F), after mitotic entry all three proteins were robustly recruited to all (100±0.0% for each MDC1, TOPBP1, and CIP2A) *γ*H2AX-containing micronuclei-derived chromatin (**Fig. 2*A-D***). Live-cell imaging of MDC1 or TOPBP1 fluorescently tagged with GFP or the Clover variant of it (to produce ^GFP^MDC1 or TOPBP1^Clover^, respectively) confirmed localization of both to damaged micronuclear chromosomes in mitosis, with levels of MDC1 and TOPBP1 on the broken micronuclear chromosome increasing more than 10-fold concomitant with chromosome condensation at mitotic entry (**Fig. 2*E, F*; Sup. Video 1 & 2**).

**Fig. 2.**
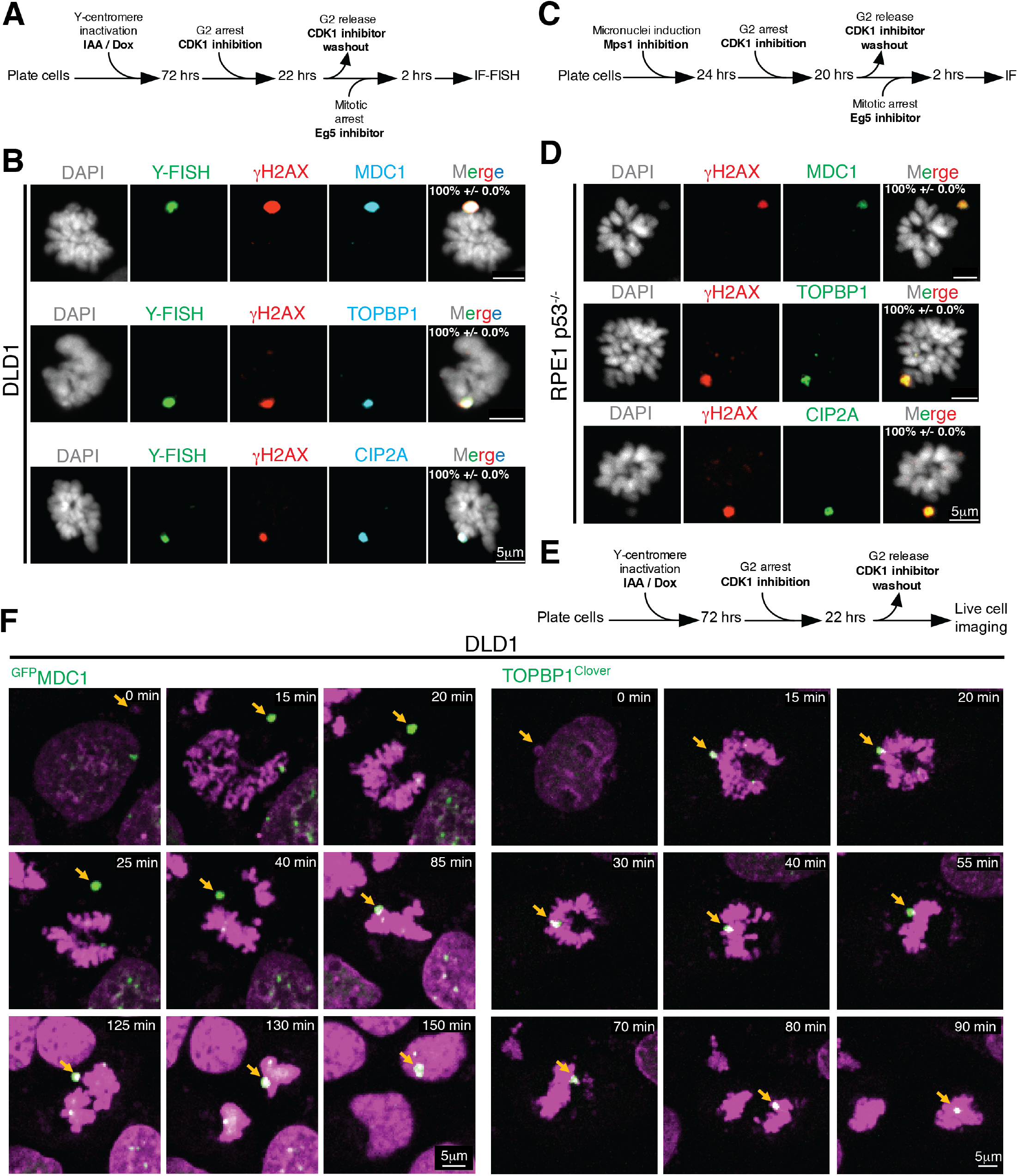
TOPBP1, MDC1, and CIP2A are recruited to a damaged micronuclear chromosome upon mitotic entry. ***(A)*** Experimental outline for (B). ***(B)*** Representative images showing localization of MDC1, TOPBP1, and CIP2A on broken micronuclear Y-chromosome in mitosis (n=3 independent experiments, total 258, 252, and 263 DLD1 cells were analyzed for MDC1, TOPBP1, and CIP2A, respectively). ***(C)*** Experimental outline for *(D)*. ***(D)*** Representative images showing localization of MDC1, TOPBP1, and CIP2A on broken micronuclear chromosome in mitosis (n=3 independent experiments, total 273, 265, and 271 RPE1 p53^-/-^ cells were analyzed for MDC1, TOPBP1, and CIP2A, respectively). Mean percentage +/- standard error for localization of the indicated protein to fragmented (*γ*H2AX positive) chromosome is indicated on the images in *(B)* and *(D)*. ***(E)*** Schematics of experimental setup for *(F)*. ***(F)*** Cropped frames of **Sup. Videos 1 and 2** showing localization of ^GFP^MDC1 and TOPBP1^Clover^ to the micronuclear chromosome during mitosis.

Damaged micronuclear chromosomes were identified to undergo one of three fates while transitioning through mitosis. In the first ∼40% of the total damaged micronuclear chromosome fragments remained clustered throughout mitosis and were incorporated into one of the daughter nuclei after cytokinesis (**Fig. 2*E, F*; Sup. Video 1 & 2**). A similar proportion of damaged and clustered micronuclear chromosome masses remained physically separated from the main chromatin mass during anaphase leading to encapsulation into a new micronucleus (**Extended Data Fig. 5*A*; Sup. Video 3**). In the final 18% of examples, damaged micronuclear chromosome fragments were clustered initially, but broke subsequently into smaller clusters from what appeared to be shear induced by mitotic spindle forces (**Extended Data Fig. 5*B*; Sup. Video 4**), a scenario likely responsible for the corresponding proportion of mitotic cells viewed in fixed imaging in which micronuclear chromosome fragments were dispersed.

### TOPBP1 and CIP2A tether chromosomal fragments in mitosis

The localization pattern of MDC1, TOPBP1, and CIP2A suggests that these proteins can mediate the tethering of fragmented micronuclear chromosomal pieces during mitosis. We tested this hypothesis by inducing Y-chromosome missegregation and then depleting MDC1, TOPBP1, or CIP2A using siRNA. Measurement of the percentage of cells with dispersed Y-fragments (**Fig. 3*A*; Extended Data Fig. 6*A***) demonstrated that depletion of any of the three (MDC1, TOPBP1, or CIP2A) resulted in the dispersal of shattered micronuclear chromosomal fragments in mitosis (**Fig. 3*B, C***). The increase in cells with dispersed Y fragments was not due to an increase in damaged Y-chromosomes as there was no statistically significant change in the frequency of cells with damaged Y-chromosome in mitosis (**Extended Data Fig. 6*B***). Thus, MDC1, TOPBP1, and CIP2A each play a role in tethering chromosomal pieces during mitosis.

**Fig. 3.**
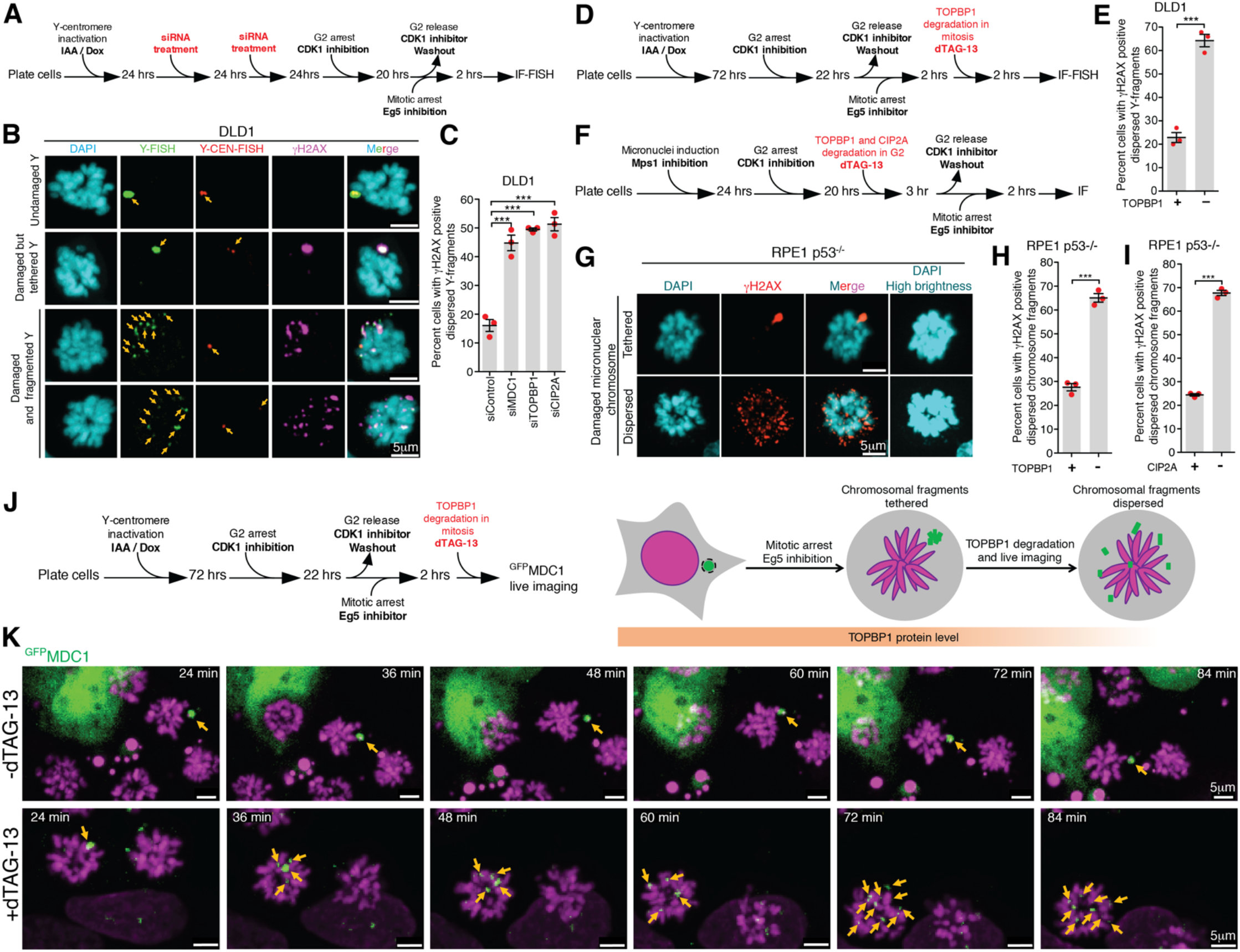
MDC1, TOPBP1, and CIP2A mediate clustering of micronuclear chromosome fragments during mitosis. ***(A)*** Experimental outline for *(B)* and *(C)*. ***(B)*** Representative images showing various fates of a micronucleated Y-chromosome in the subsequent mitosis. ***(C)*** Quantitation of cells with dispersed Y-chromosome fragments in mitosis upon siRNA mediated depletion of MDC1, TOPBP1, and CIP2A (n=3 independent experiments; a total of 770, 686, 662, and 667 cells were analyzed for control, siMDC1, siTOPBP1, and siCIP2A conditions, respectively). (One-way analysis of variance with Bonferroni’s multiple comparison test was applied, *** P<0.0001 and ns P>0.05.) ***(D)*** Schematic of experimental outline for *(E)*. ***(E)*** Quantitation of cells with dispersed Y-chromosome fragments in mitosis upon degradation of TOPBP1^dTag^ in mitosis (n=3 independent experiments; total 997 and 1131 cells were analyzed for control and TOPBP1^dTag^ degradation condition, respectively). ***(F)*** Experimental outline for *(G-I)*. ***(G)*** Representative images of RPE p53^-/-^ cells showing tethered and dispersed damaged micronuclear fragments in mitosis after induction of micronucleation by Mps1 inhibition. Quantitation of cells with dispersed micronuclear fragments in mitosis upon degradation of ***(H)*** TOPBP1^dTag^ (n=3 independent experiments; total 476 and 688 cells were analyzed for control and TOPBP1^dTag^ degradation condition, respectively) or ***(I)*** ^dTag^CIP2A (n=3 independent experiments; total 511 and 445 cells were analyzed for control and ^dTag^CIP2A degradation conditions respectively). ***(J)*** Experimental outline and schematic for *(K)*. ***(K)*** Frames of **Sup. Video 5 and 6** showing behavior of ^GFP^MDC1 labelled clustered damaged micronuclear chromosome in mitosis, when TOPBP1^dTag^ degradation is induced (bottom) or not (top). Yellow arrows in *(B)* and *(K)* point to the fragments of micronuclear chromosomes. Two-tailed unpaired t-test was applied for data shown in *(E), (H)*, and *(I)*; *** P<0.0001. For graphs in *(C), (E), (H)* and *(I)*, mean and standard error are shown, individual data points are shown as solid red circles.

TOPBP1 plays multiple distinct roles throughout the cell cycle^31^. To test if TOPBP1 acts as a protein tether for chromosomal fragments in mitosis or if the tethering of chromosomal pieces during mitosis is a consequence of its activity in the earlier cell cycle phase, we perturbed TOPBP1 levels at specific phases of the cell cycle. To achieve this, we used CRISPR mediated genome editing to modify both endogenous TOPBP1 loci to encode a variant (TOPBP1^dTag^) containing a carboxyterminal degradation tag (dTag)^32^ (**Extended Data Fig. 7*A***). TOPBP1^dTag^ degradation was then induced by addition of the heterobifunctional small molecule dTAG-13 to mediate binding of the dTag to the E3 ubiquitin ligase cereblon (CRBN), producing rapid and efficient degradation of TOPBP1^dTag^ with a half-life of 14 min (**Extended Data Fig. 7*C, D***).

We tested if the S-phase role(s) of TOPBP1 in regulating DNA replication and ATR activation^31,33^ is sufficient for micronuclear fragment clustering in mitosis. We induced Y-centromere inactivation and micronucleation and arrested cells in the subsequent G2 phase (using the CDK1 inhibitor RO-3306). We then induced TOPBP1 degradation in G2 and released the cells into mitosis by removal of the CDK1 inhibitor (**Extended Data Fig. 7*F***). Depletion of TOPBP1 in the G2 phase resulted in a three-fold increase in cells with dispersed Y fragments in mitosis (**Extended Data Fig. 7*G***). Despite TOPBP1 depletion in G2, we did not observe any change in the percent of mitotic cells with a damaged Y chromosome, indicating that the DNA damage or repair frequency was not altered (**Extended Data Fig. 7*H***). We conclude that beyond any G1 or S-phase roles of TOPBP1, tethering of chromosomal pieces in mitosis requires continued presence of TOPBP1 in G2.

TOPBP1 regulates mitotic DNA synthesis and the recruitment of various DNA repair proteins to some *γ*H2AX positive chromatin sites during early mitosis^34-36^. We tested if chromosome fragment tethering is a consequence of repair events in early mitosis or if TOPBP1 acts as a protein tether for chromosomal fragments throughout mitosis. To do this, we induced Y-chromosome micronucleation and arrested cells in the subsequent mitosis by releasing G2 arrested cells for 2 hours into media containing the Eg5 inhibitor. Induction of TOPBP1^dTag^ loss in the resultant mitotically arrested cells (**Fig. 3*D***) produced a three-fold increase in the percentage of cells with dispersed of micronuclear fragments without any change in the proportion of mitotic cells with a damaged Y chromosome (**Extended Data Fig. 7 *E, I***).

We next followed the behavior during mitosis of clustered Y chromosomal fragments during the induced degradation of TOPBP1^dTag^. Since MDC1 is known to recruit TOPBP1 to damaged chromosomes during mitosis but not vice versa^27^, we used a GFP-tagged MDC1 (^GFP^MDC1) to mark damaged micronuclear chromosomes. The ability of ^GFP^MDC1 to localize to the damaged chromosomes independent of TOPBP1 makes it an ideal marker to follow the behavior of micronuclear chromosome fragments throughout mitosis in the presence or absence of TOPBP1 (**Fig. 3*J***). Live cell imaging revealed that in mitotically arrested cells all ^GFP^MDC1 positive clusters of chromosomal fragments remained tethered for the full 2-hour span of filming (**Fig. *3K*, top panels; Sup. Video 5**). However, when TOPBP1^dTag^ degradation was induced during imaging, ^GFP^MDC1-bound chromosome fragments dispersed within 30 minutes after the addition of dTAG-13 (**Fig. 3*K*, bottom panels; Sup. Video 6**). Thus, beyond possible roles in G1, S and G2 phases, continuing presence of TOPBP1 is essential for maintaining micronuclear chromosome fragment clustering in mitosis.

Next, we used CRISPR mediated genome editing to modify both endogenous CIP2A loci of RPE p53^-/-^ cells to encode ^dTag^CIP2A. After induction of micronuclei or chromatin bridge formation (produced by Mps1 or Topoisomerase II inhibition, respectively), induced degradation in the subsequent G2 phase of ^dTag^CIP2A led to loss of tethering of chromosome fragments marked by *γ*H2AX (**Fig. 3*F, G*, and *I*; Extended Data Fig. 7 *B*; Extended Data Fig. 8*A-D***). As its name suggests, CIP2A was identified as an inhibitor of PP2A^37^. We tested if CIP2A’s role in clustering of micronuclear chromosomal fragments in mitosis is through its inhibition of PP2A. We induced micronuclei formation (by Mps1 inhibition) and arrested the cells in the subsequent G2 phase. We then inhibited PP2A activity for 1 hour using okadaic acid, induced degradation of ^dTag^CIP2A, and finally released the cells into mitosis (**Extended Data Fig. 9*A***). The percentage of cells with dispersed micronuclear chromosomal fragments after CIP2A degradation was unaffected by PP2A inhibition (**Extended Data Fig. 9*B, C***). We thus conclude that tethering of chromosomal fragments in mitosis generated as a result of either micronucleation or chromatin bridge formation requires CIP2A in and that PP2A inhibition is not the only role that CIP2A plays in mediating and maintaining clustering of micronuclear chromosome fragments during mitosis.

### Dispersal of chromosomal fragments in mitosis leads to their random segregation between daughter cells

To test the consequence of disrupting chromosomal fragment clustering during mitosis (**Fig. 4*A*)**, we induced chromosome missegregation and micronucleation by Mps1 inhibition in RPE1 p53^-/-^ cells and monitored the distribution of broken chromosome fragments using *γ*H2AX in early G1 or telophase (**Fig. 4*B***). Aurora-B staining at the midbody was used to identify paired daughter cell nuclei. Disruption of fragment clustering by degradation of TOPBP1^dTag^ or ^dTag^CIP2A resulted in a random distribution of chromosomal fragments between the resulting daughter cells in a large majority (69.8±0.15% and 70.7±1.9% for TOPBP1^dTag^ and ^dTag^CIP2A degradation condition, respectively) of cases with some chromosomal fragments encapsulated within a main nucleus and others in the cytoplasm of daughter cells (**Fig. 4*C, D***).

**Fig. 4.**
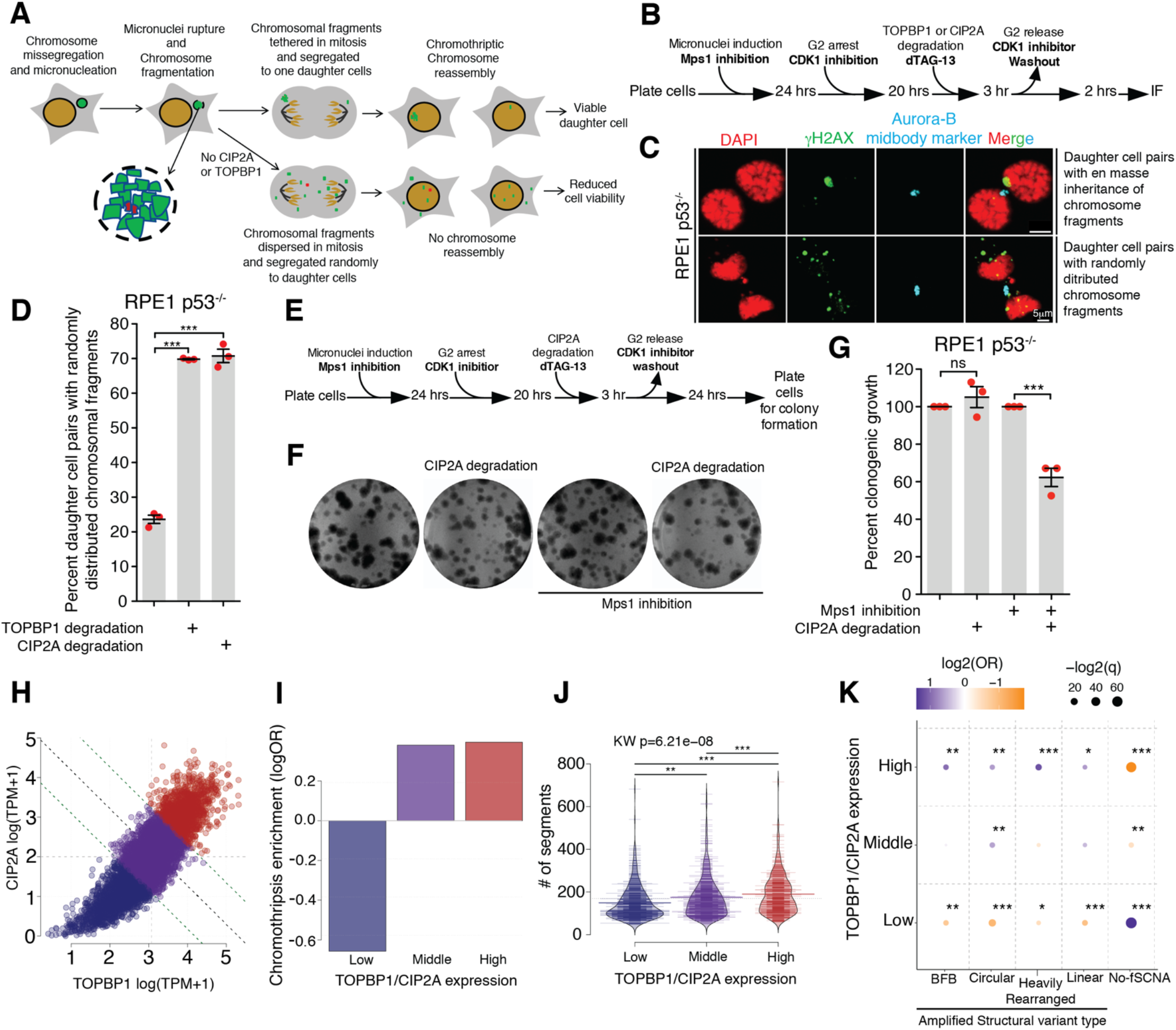
TOPBP1 and CIP2A are enriched in chromothriptic tumor samples and allow transfer of fragments to the same daughter cell essential for viability of the resulting daughter cell. ***(A)*** Schematic showing sequence of events and fates of daughter cell post micronucleation in indicated conditions. ***(B)*** Experimental outline for *(C)* and *(D)*. ***(C)*** Representative images showing various patterns of segregation of damaged micronuclear chromosome fragments in telophase or G1 phase. ***(D)*** Quantitation of cells with dispersed micronuclear fragments distributed between daughter cells following micronuclei formation and induced degradation in G2 of TOPBP1^dTag^ or ^dTag^CIP2A (n=3 independent experiments; total 306, 311, and 339 cells were analyzed for control, TOPBP1^dTag^ degradation, and ^dTag^CIP2A degradation conditions, respectively). (One-way analysis of variance with Bonferroni’s multiple comparison test was applied, *** P<0.0001.) ***(E)*** Experimental outline for *(F)* and *(G)*. ***(F)*** Images of colony formed for experiment outlined in *(E)* for indicated conditions. ***(G)*** Quantitation of number of colonies formed for the experiment outlined in *(E)* for the indicated conditions (n=3 independent experiments; One-way analysis of variance with Bonferroni’s multiple comparison test was applied, *** P<0.0001 and ns P>0.05). ***(H)*** Expression levels (log(TPM+1)) of TOPBP1 and CIP2A in 7,856 tumor samples from The Cancer Genome Atlas (TCGA). Samples are categorized as joint upper (red), middle (purple) or lower (blue) expression of both genes by bisecting the line of best fit with a decision boundary at the upper quartile or lower quartiles of each gene (green dotted lines). Light grey dotted lines denote median values of gene expression, black dotted line indicates the bisecting line of the median. TPM=transcripts per million. ***(I)*** Enrichment of chromothriptic samples in the gene expression groups defined in (*H)*. Two-sided Fisher’s exact test on contingency tables of chromothriptic/non-chromothriptic and gene expression group vs. rest. All q<0.001. OR=odds ratio. ***(J)*** Association between the number of copy number segments in a copy number profile (y-axis) and gene expression groups defined in *(H)*. Thick colored horizontal lines indicate mean number of segments for each expression group. Thin horizontal lines indicate individual samples. Black horizontal line indicates the mean number of segments across the dataset. Overall significance was tested with a Kruskal-Wallis test. Pairwise comparisons were performed with two-sided Mann-Whitney tests. **p<0.01, ***p<0.001. KW=Kruskal-Wallis test. ***(K)*** Associations between gene expression groups defined in *(H)* (y-axis), and classes of amplicon events defined by AmpliconArchitect^41^ (x-axis), using two-sided Fisher’s exact tests. Correction for multiple testing was performed using the Benjamini-Hochberg method. Color denotes odds ratio, size of points indicates the significance of the test. *q<0.05, **q<0.01, ***q<0.001. BFB=breakage fusion bridge event. No-fSCNA=no focal somatic copy number amplification. OR=odds ratio.

### Clustering of micronuclear chromosomal fragments during mitosis is required for the viability of the daughter cells

Viability of cells undergoing chromosomal instability and forming frequent micronuclei will require continued expression in daughter cells of all essential genes (**Fig. 4*A***), while dispersal of fragments and the loss of essential genes/chromosomal regions (including centromeres or telomeres) would result in inviable daughter cells. We tested how viability was affected by the absence of mitotic clustering to enable delivery of most or all of them to the same daughter nucleus for their subsequent ligation into a highly rearranged, heritable chromosome. We induced micronuclei formation in p53^-/-^, CIP2A^dTag^-expressing RPE1 cells (by treating them for 24 hours with an Mps1 inhibitor) and then induced degradation of CIP2A^dTag^ in the subsequent cell-cycle (**Fig. 4*E***). CIP2A depletion reduced cell viability by 2-fold, as measured with a colony formation assay (**Fig. 4*F, G***).

### Abundant chromothripsis in tumors retaining high levels of Cip2A, TOPBP1, and MDC1

The impact of chromosomal fragment tethering on the cancer genome was first assessed by using 7,856 samples from The Cancer Genome Atlas (TCGA) project where cancer RNA expression data was associated with chromothripsis events identified using CTLPscanner, which evaluates the oscillation of copy number changes measured using single nucleotide polymorphism (SNP) arrays^38^. An additional analysis was also performed using whole genome sequencing-derived chromothripsis events (determined with ShatterSeek^39^) in the 667 tumors from the Pan Cancer Analysis of Whole Genome (PCAWG) project for which gene expression data was available. Accumulation of RNAs encoding TOPBP1, CIP2A, and MDC1 was highly correlated with each other, with each more highly expressed in samples in which chromothripsis was identified by copy number (change in mean log TPM+1=0.15, 0.20, 0.16, and p=9.3e^-57^, 2.4e^-14^, 1,1e^-16^ for *TOPBP1, CIP2A* and *MDC1* respectively, two-sided Mann-Whitney tests) or structural variants (change in mean log TPM+1=0.22, 0.44, 0.28, and p=0.031, 3.0e^-4^, 7.7e^-3^ for *TOPBP1, CIP2A* and *MDC1* respectively, two-sided Mann-Whitney tests; **Extended Data Fig. 10*A-D***).

We further categorized samples into low, middle, or high expression of CIP2A and TOPBP1 by bisecting the line of best fit at the upper and lower quartiles (**Fig. 4*H***). This categorization confirmed an enrichment of chromothripsis in highly expressing samples (OR=0.39, p=6.2e^-4^, two-sided Fisher’s exact test), and a depletion in low expressing samples (OR=-0.66, p=1.9e^-12^, two-sided Fisher’s exact test; **Fig. 4*I***). Moreover, our analysis revealed positive associations of CIP2A and TOPBP1 with copy number variations, segmentation of copy number profile, and extent of loss of heterozygosity (LOH) expected from chromothripsis in both datasets (**Fig. 4*J***; **Extended Data Fig. 11*A-C***). In low TOPBP1/CIP2A expressing chromothriptic samples in TCGA, there was a significant depletion (OR=0.48, q=3.2e^-4^) of chromothripsis-amplification type copy number signatures^40^ and a trend (OR=1.5, p = 0.14) towards enrichment in high expressing samples (**Extended Data Fig. 11*D***), an outcome confirmed by enrichment of multiple classes of amplicon structures^41^ only in high expressing samples of both non- and chromothriptic designation (**Fig. 4*K***).

## Discussion

Here we identify that a complex of TOPBP1 and CIP2A recruited by MDC1 mediates the tethering of chromosomal fragments in mitosis generated as a result of either micronucleation or chromatin bridge formation. Such tethering is a crucial step in chromothriptically produced genome rearrangements, the *en masse* inheritance of most or all micronuclear chromosome fragments by one of the daughter cells. Initial chromosome fragmentation within damaged or ruptured interphase micronuclei recruits megabase domains of *γ*H2AX that rapidly recruit MDC1 released by disassembly of the main nuclear envelope. In turn, MDC1 assembled at or near each DSB directly recruits TOPBP1 and its binding partner CIP2A. The assemblies of TOPBP1 and CIP2A are essential for tethering of chromosomal fragments throughout mitosis and facilitate their incorporation into the primary nucleus of one daughter cell where they remain in close proximity, facilitating their ligation by NHEJ into a heritable chromothriptic chromosome (**Extended Data Fig. 12**). The nuclear foci resulting from inheritance of the clustered micronuclei fragments in the daughter nuclei may be similar to the recently described micronuclear body^42^. While the absence of rounded droplets in live cell imaging offers no support for chromosome fragment tethering through a mechanism involving TOPBP1 condensation^43^, chromosome fragment clustering throughout mitosis may be mediated by the multiple phosphate-binding BRCT domains of TOPBP1^44^ and/or by CIP2A oligomerization through its proposed coiled-coil domain.

While the mitotic clustering of the micronuclear chromosome fragments may suggest minimal loss of DNA copy during chromothripsis, the chromothriptic signature in cancer genomes most often shows oscillation of copy number including loss of some DNA fragments^45^. Despite MDC1/TOPBP1/CIP2A tethering, losses may be mediated by multiple mechanisms, including *(i)* selective pressures, including the loss of tumor suppressor genes, *(ii)* spindle forces breaking initially clustered micronuclear chromosome fragments into smaller groups (as we observed in about 20% of cases in our live-cell imaging - **Extended Data Fig. 5*B***), and *(iii)* altered gene copy numbers from defective DNA replication in micronuclei^3,9^ or exo-nuclease attack on the micronuclear DNA either in interphase^5,6^ or mitosis. We have shown that chromosome fragment clustering in mitosis affects viability of the resultant daughter cells (**Fig. 4*E-G***). Consistent with this, our analysis of cancer genomic data has identified that tumors with a chromothriptic signature retain high level expression of MDC1, TOPBP1, and CIP2A. Since CIP2A is non-essential to dividing cells and its expression is low in most normal human tissues but high in many cancers^46^, we propose that it is an attractive therapeutic target for chromosomally unstable tumors forming frequent micronuclei.

## Materials and Methods Cell culture

HEK293T (ATCC: CRL-11268), DLD-1, and hTERT-RPE1 p53^-/-^ (a gift kind from Prof. David Pellman) cells were cultured in Dulbecco’s modified Eagle’s medium (DMEM, Gibco) or DMEM/F-12 (Gibco) (for hTERT-RPE1 p53^-/-^) media, supplemented with 10% tetracyclin-free fetal bovine serum (FBS), 2mM L-Glutamine, and 100ug/ml Normocin (Invivogen) at 37°C in presence of 5% CO_2_. Cell lines were periodically tested and confirmed free of mycoplasma.

### Stable cell line generation and genome editing

For lentivirus production TOPBP1-clover cloned into LS135 or lenti-CMV/TO-GFP-MDC1 (779-2) (a gift from Eric Campeau, Addgene plasmid # 26285) were co-transfected with packaging plasmids pMD2.G and psPAX2 in HEK293T using TransIT-VirusGEN® Transfection Reagent (Mirus, MIR6705). The supernatant containing lentivirus was collected and filtered through a 0.45μm filter, 2-3 days after transfection. For generating stable cell lines expressing fluorescently tagged proteins, cells were infected with viral supernatant in presence of 5g/ml Polybrene (Santa Cruz). Single-cell clones expressing the protein of interest were isolated by fluorescence-activated cell sorting (FACS) (Sony SH800).

For tagging endogenous locus with dTAG, sgRNA sequences targeting the desired genomic region (shown in **Extended Data Fig. 7*A, B***) were cloned in pSpCas9(BB)-2A-GFP (PX458) (a gift from Feng Zhang, Addgene plasmid # 48138). The repair template was created by cloning the left and right homology arm (∼500bp) with 2xHA-dTAG (for N-term tagging) or dTAG-2xHA (for N-term tagging) into pUC19. Cas9 and sgRNA expressing plasmid (PX458) along with plasmid containing repair template were nucleofected in DLD1 or hTERT-RPE1 p53-/- using Cell Line Nucleofector™ Kit V (Lonza) and Nucleofector® 2b Device (Lonza) according to manufacturer’s protocol. Single cells expressing GFP were FACS sorted (Sony SH800) 2 days post nucleofection and grown to colonies. PCR and Immunoblotting were used to screen colonies derived from single cells to identify correctly edited colonies.

Following sgRNAs were used for editing and PCR primers were used for screening colonies: sgTOPBP1: AGATGCGATTAGTGTACTCT

sgCIP2A: GGCAGTGGAGTCCATTGCAC

PCR primers for confirming editing at TOPBP1 locus:

For DLD1 cells:

TOPBP1fw-1: GGTGAGACTTTGTCCCACAGGGT

TOPBP1rv-1: AACCTTGTGCTCAGGCTCCTGTT

For RPE1 p53^-/-^ cells:

TOPBP1fw-2: GAGACTTTGTCCCACAGGGTCCA

TOPBP1rv-2: AACCTTGTGCTCAGGCTCCTGTTA

PCR primers for confirming editing at CIP2A locus:

CIP2Afw: ATGCAGGCTCTGGCGGAGTG

CIP2Arv: ACGCTTACTAGGAAGGGGAAGTGC\

### siRNA transfection

ON-TARGETplus SMARTpool siRNAs (Horizon Discovery) targeting either Lig4 (L-004254-00-0005), PolD3 (L-026692-01-0005), Mre11 (L-009271-00-0005), Rad50 (L-005232-00-0005), PolQ (L-015180-01-0005), MDC1 (L-003506-00-0005), TOPBP1 (L-012358-00-0005), CIP2A (L-014135-001-0005) or non-targeting pool (D-001810-10-05) were transfected using Lipofectamine RNAiMAX Transfection Reagent (ThermoFisher, 13778075) at the indicated timepoint according to manufacturer’s protocol.

### Inhibitors and small molecules

Doxycycline was used at 1 μg/ml (Sigma), Indole-3-acetic acid (IAA) used at 500 μM (Sigma, I5148), CDK1 inhibitor RO-3306 used at 9 μM (Sigma-Aldrich, SML0569), NMS-P715 0.8 μM (Millipore Sigma, 475949), ICRF-193 used at 100 nM (Enzo life sciences, BML-GR332-0001) S-Trityl-L-cysteine (STLC) used at 5 μM (Enzo Life Sciences, ALX-105-011-M500), dTAG-13 used at 500 nM (Sigma-Aldrich, SML2601), Okadaic acid used at 500 nM (Fisher Scientific, 11-362-5U), Geneticin® Selective Antibiotic (G418 Sulfate) used at 300 μg/ml (Gibco, 10131035).

### Immunoblot (IB)

Cells were lysed in SDS sample buffer and boiled at 95°C for 10 minutes. The whole cell lysate was resolved on SDS PAGE gel and transferred to Immobilon®-FL PVDF membrane (Millipore Sigma). The membrane was then blocked with Intercept® (TBS) Blocking Buffer (LI-COR) for 1 hour followed by overnight incubation at 4°C with primary antibody diluted in Intercept® (TBS) Blocking Buffer supplemented with 0.1% Tween-20. The membrane was washed 4 times (5min each) with TBST (TBS, 0.1% Tween-20) and then incubated for 1 hour at room temperature with fluorescent secondary antibody diluted in Intercept® (TBS) Blocking Buffer supplemented with 0.1% Tween-20. The membrane was washed 4 times (5 minutes each) with TBST (TBS, 0.1% Tween-20) and then imaged on Odyssey FC imager (LI-COR).

### Immunofluorescence (IF)

Cells were plated on coverslips and treated as indicated. Cells were fixed with 4% PFA in PBS or 100% ice-cold methanol (for TOPBP1 staining) for 15 minutes at room temperature, followed by permeabilization with 0.5% Triton X-100 in PBS for 15 minutes at room temperature. Cells were then blocked with blocking buffer (0.2M glycine, 2.5% BSA, 0.1% Triton X-100, PBS) for 1 hour at room temperature. Coverslips were incubated with primary antibody diluted in blocking buffer for 1 hour at room temperature or overnight at 4°C. Coverslips were washed 4 times with PBST (PBS, 0.1% Tween-20), 10 minutes each at room temperature, followed by incubation with fluorescent secondary antibodies diluted in blocking buffer for 1 hour at room temperature. Coverslips were washed 4 times with PBST (PBS, 0.1% Tween-20), 10 minutes each at room temperature. Coverslips were then either processed for Fluorescent In-Situ Hybridization (FISH) or stained with DAPI and mounted on slides using Prolong Gold mounting media. For combined IF and FISH, cells were fixed with methanol/acetic acid (3:1) after IF procedure and then processed for FISH as described below.

### Antibodies

Following antibodies were used in this study at the indicated concentration. Anti-phospho histone H2AX (Ser139) (*γ*H2AX) (clone JBW301, EMD Millipore, 05-636, 1:1500 IF) (Cell Signaling Technologies, 2577, 1:1000 IF), anti-TOPBP1 (Bethyl Laboratories, A300-111A, 1:2000 IF, 1:2000 IB) (Abcam, ab2402, 1:1000 IF), anti-MDC1 (Abcam ab11171, 1:1000 IF, 1:1000 IB), anti-CIP2A (clone 2G10-3B5; Santa Cruz sc80659, 1:500 IF, 1:1000 IB), anti-GAPDH (clone 14C10, Cell Signaling Technologies, 1:5000 IB), anti-PolD3 (clone 3E2, Abnova, H00010714-M01, 1:1000 IB), anti-Lig4 (clone N2C2, GeneTex, GTX100100, 1:1000 IB), anti-Mre11 (clone 12D7, GeneTex, GTX70212, 1:1000), anti-Rad50 (Cell Signaling Technologies, 3427, 1:1000 IB), anti-HA-tag (Novus biologicals, NB600-363, 1:2000 IB), anti-Lamin A/C (clone E-1, Santa Cruz, sc-376248, 1:200 IF), anti-Lamin B1 (clone C-5, Santa Cruz, sc-365962, 1:200 IF), anti-53BP1 (Thermo Fisher Scientific, PA1-16565, 1:1000 IF), anti-BRCA1 (clone D-9, Santa Cruz, sc-6954, 1:1000 IF).

### Chromosome spreads and Fluorescent In-Situ Hybridization (FISH)

For metaphase chromosome spreads, cells were arrested in mitosis for 4 hours using 100ng/ml colcemid (KaryoMAX, Thermo Fisher). The cells were then collected by trypsinization and incubated with hypotonic solution (25 mM KCl, 0.27% sodium citrate in distilled water) for 10-15 minutes at 37°C followed by fixation with cold methanol/acetic acid (3:1). Fixed cells were then pelleted and resuspended in appropriate concentration in fixative and dropped on the slides for chromosome spread preparation.

For DNA FISH, human Y-chromosome and human Y-centromere FISH probes (MetaSystems) were combined in a 1:1 ratio and applied to chromosome spreads and covered with coverslip or applied to cells on coverslips (for combined IF-FISH) followed by denaturation at 75°C for 5 minutes. The coverslips were sealed with Fixogum rubber cement (Marabu) and incubated overnight in a humidified chamber at 37°C for hybridization. Slides or coverslips were then washed first with 0.4% SSC buffer at 72°C for 2 minutes, and then by 2X SSC with 0.05% Tween-20 for 30 seconds at room temperature. Slides or coverslips were then washed with distilled water and counterstained with DAPI and mounted in Prolong Gold mounting media.

### Fixed cell and Live cell Imaging

Fixed cell images were acquired on the DeltaVision Core system (Applied Precision) and maximum intensity projections were generated using the Fiji package of ImageJ. Fixed images were also acquired on the CQ1 benchtop spinning disk confocal high-content analysis system (Yokogawa).

For live-cell imaging, DLD1 or RPE1 p53^-/-^ expressing the fluorescently tagged protein of interest were plated in 96 well glass-like polymer bottom plates (Cellvis) and treated as indicated. Live cell images were acquired for 4-24 hours CQ1 benchtop spinning disk confocal high-content analysis system (Yokogawa). Cells were imaged at 40X or 60X dry objective and were maintained in humidified condition at 37°C in presence of 5% CO_2_ during imaging.

### Clonogenic assay

Cells were treated as indicated in the experimental outline and the specified number of cells were plated. For the experiment shown in the **Extended Data Fig. 2*G-J***, cells were grown in 300 μg/ml G418 containing media. After 10-15 days the cells were fixed with 100% methanol and stained with a solution of 0.5% crystal violet, 25% methanol. The colonies formed were manually quantified.

### Statistical analysis

Three independent replicates were performed for each experiment unless otherwise stated. Prism v5.0 (GraphPad) was used for statistical analysis except for **Fig. 4*H-K*** and **Extended Data Fig. 10 and 11**. All graphs show mean and standard error with individual data points for each repeat. The statistical test used for analysis is indicated in figure legends.

### Gene expression analysis

Gene expression data (gene level counts and TPMs) for TCGA were downloaded from the GDC portal (https://portal.gdc.cancer.gov). Processed copy number profiles from SNP6 arrays for TCGA were obtained from: https://github.com/VanLoo-lab/ascat/tree/master/ReleasedData/TCGA_SNP6_hg19. Chromothripsis calls for PCAWG (ShatterSeek) were downloaded from: https://dcc.icgc.org/releases/PCAWG/evolution_and_heterogeneity/clustered_mut_processes.

Chromothripsis was called from SNP6 copy number profiles using CTLPscanner^47^. Associations between log(TPM+1) *CIP2A, TOPBP1*, and *MDC1* gene expression and chromothripsis were performed with two-sided Mann-Whitney tests between chromothriptic and non-chromothriptic samples. This was performed separately for chromothripsis defined by CTLPscanner (TCGA SNP6) and chromothripsis defined by ShatterSeek (PCAWG WGS). Expression of *CIP2A* and *TOPBP1* was jointly categorized into lower, middle and upper by finding perpendicular lines to the line of best fit that pass through both the upper quartile and lower quartile for both *TOPBP1* and *CIP2A*. In brief the line of best fit for log(TPM+1) gene expression of *TOPBP1* and *CIP2A* is given as

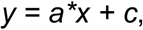

where *y* is the gene expression for *CIP2A, x* is the gene expression for *TOPBP1, a* is the gradient and *c* is the intercept. A perpendicular line that passes through a point defined as [xi,yi] is then defined with coefficients:

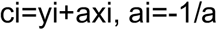

Where [xi,yi] are given as the upper quartiles of *TOPBP1* and *CIP2A* expression to determine a threshold for upper expression, and given as the lower quartiles of *TOPBP1* and *CIP2A* expression to determine a threshold for lower expression.

Associations between categorized *CIP2A/TOPBP1* expression and chromothriptic samples were performed using two-sided Fishers exact tests. Associations between categorized *CIP2A*/*TOPBP1* expression and the number of segments in copy number profiles were tested with a Kruskal-Wallis rank-sum test, with subsequent pairwise two-sided Mann-Whitney tests. Associations between categorized *CIP2A/TOPBP1* expression and the proportion of the genome as LOH were tested with a Kruskal-Wallis rank-sum test, with subsequent pairwise two-sided Mann-Whitney tests. Categorization and subsequent associations were performed for SNP6 data in all samples, for SNP6 data in only chromothriptic samples, and for WGS data in only chromothriptic samples. Copy number signature exposures for TCGA copy number profiles were downloaded from Steele *et al*. 2022^40^. Associations between categorized *CIP2A/TOPBP1* expression and chromothripsis-associated copy number signatures (CN4:9) were performed using Fisher’s exact tests. Correction for multiple testing was performed using the Benjamini-Hochberg method. Samples categorized by their amplicon type as defined by AmpliconArchitect were downloaded from Kim et al 2020 ^41^. Associations between amplicon type and categorized *CIP2A/TOPBP1* expression were performed using Fishers exact tests. Correction for multiple testing was performed using the Benjamini-Hochberg method.

## Data availability

All data will be available upon request.

## Code availability

No custom algorithms were developed for the bioinformatic analysis. Data analysis code is available upon request.

