## Extended Data for "Mitotic tethering enables *en masse* inheritance of a shattered micronuclear chromosome"

**Trivedi et al. Extended Data Fig. 1 Fragmentation of Y-chromosome after micronucleation.**

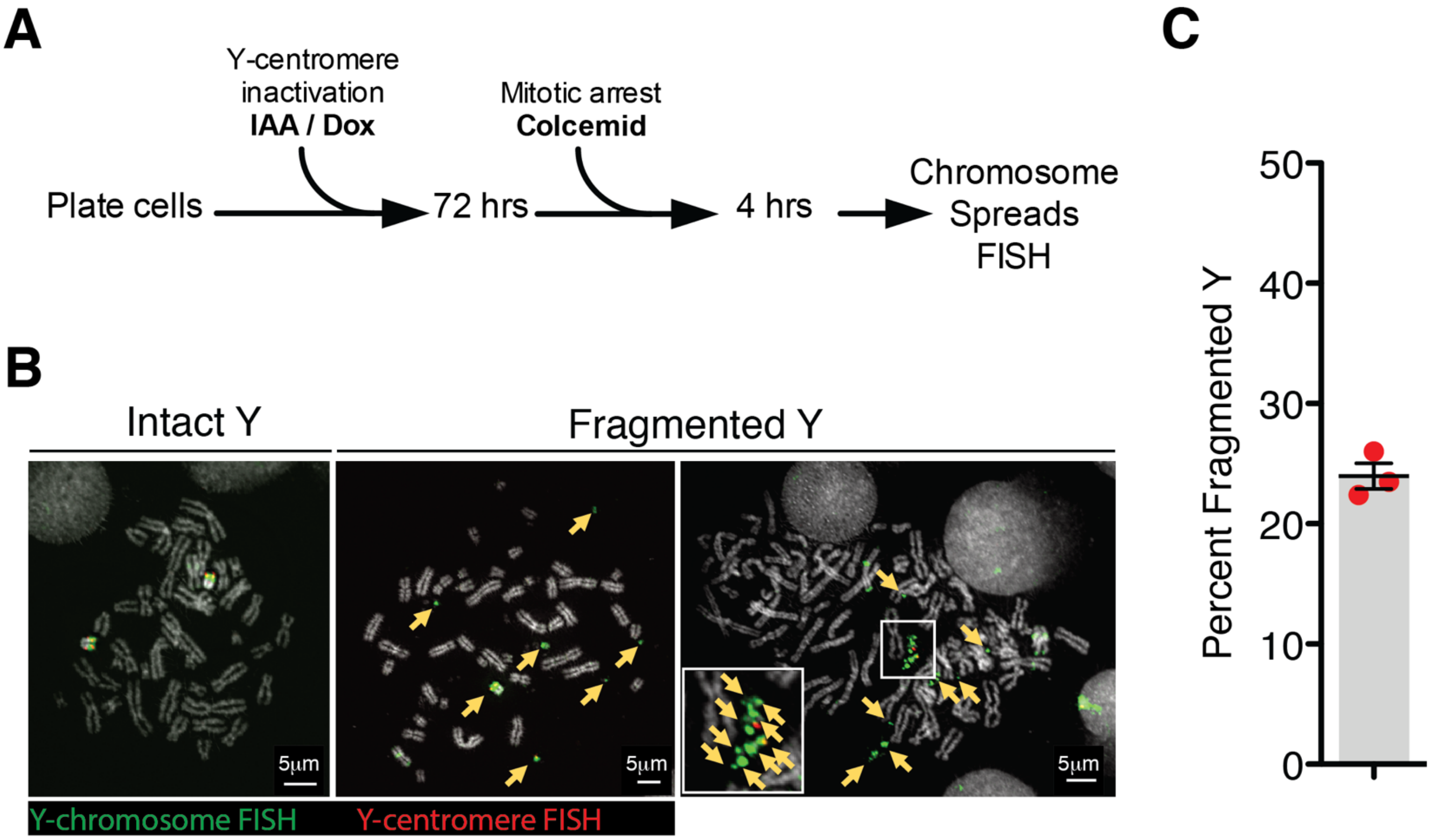


**Extended Data Fig. 1. Fragmentation of Y-chromosome after micronucleation.** ***(A)*** Experimental schematic for *(B)* and *(C)*. ***(B)*** Representative images of Y-chromosome in chromosome spreads from experiment outlined in *(A)*. Yellow arrows point to Y-chromosome fragments. ***(C)*** Quantitation of fragmented Y-chromosome from experiments outlined in *(A)* (n=3 independent experiments, total 203 cells were analyzed).

**Trivedi et al. Extended Data Fig. 2 Micronuclear envelopes do not persist in mitosis and PolD3 does not mediate clustering of broken chromosome fragments in mitosis.**

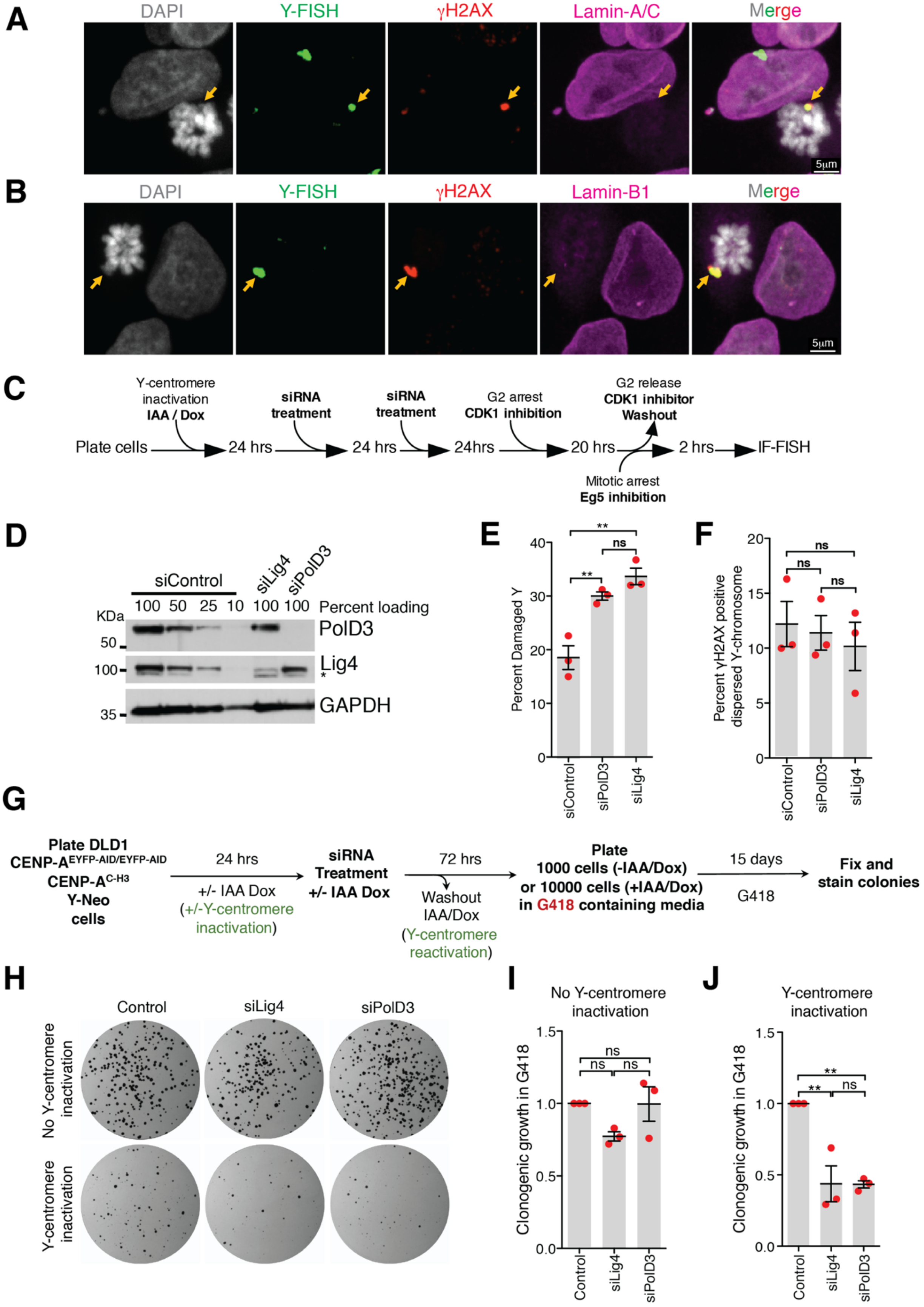


**Extended Data Fig. 2. Micronuclear envelopes do not persist in mitosis and PolD3 does not mediate**

**clustering of broken chromosome fragments in mitosis.** Representative image showing ***(A)*** Lamin A/C or ***(B)*** Lamin B1 staining on damaged and clustered Y-chromosome in mitosis. ***(C)*** Schematic for experiment in *(D)*, *(E)*, and *(F)*. ***(D)*** Immunoblots showing depletion of PolD3 and Lig4 from experiment described in *(C)*. ***(E)*** Quantitation of damaged Y-chromosome for experiment shown in *(C)*. ***(F)*** Quantitation of fragment dispersal of a damaged Y-chromosome for experiment shown in *(C)* (for *(E)* and *(F)*, n=3 independent experiments; total 651, 555, and 637 cells were analyzed for control, siPolD3, and siLig4 conditions, respectively). (One-way analysis of variance with Bonferroni's multiple comparison test was applied, ** P<0.001 and ns P>0.05.) ***(G)*** Schematic of experimental setup for *(H)*, *(I)*, and *(J)*. ***(H)*** Images of CEN-select colony formation assay from experiment outlines in *(G)*. ***(I, J)*** Quantitation of clonogenic growth from the experiment outlined in *(G)* and shown in *(H)* (for *(I)* and *(J)*, n=3 independent experiments were analyzed; one-way analysis of variance with Bonferroni's multiple comparison test was applied, ** P<0.001 and ns P>0.05).

**Trivedi et al. Extended Data Fig. 3. Neither the MRN complex nor PolQ mediates clustering of broken chromosome fragments in mitosis and micronuclei are defective in recruitment of DNA repair proteins.**

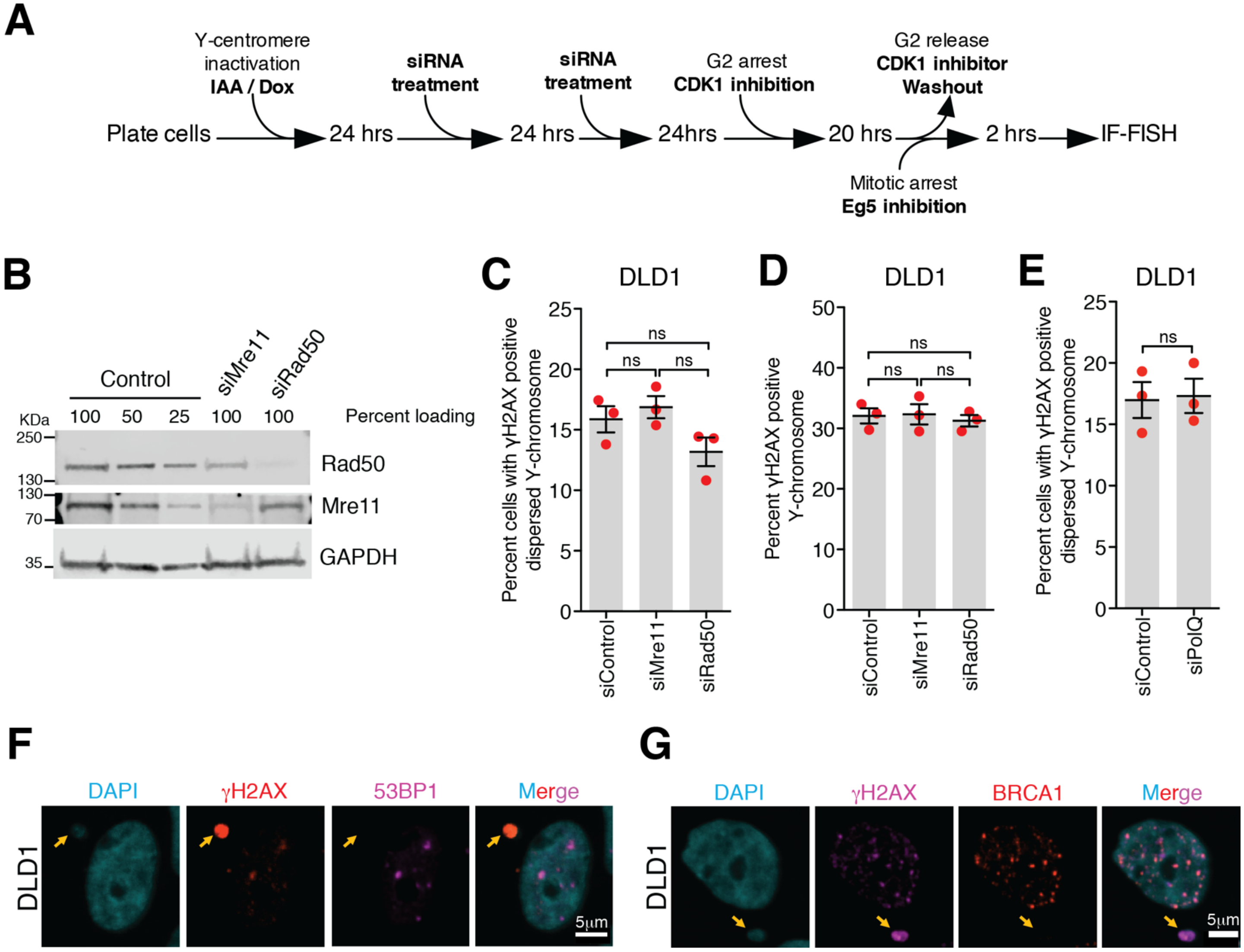


**Extended Data Fig. 3. Neither the MRN complex nor PolQ mediates clustering of broken chromosome fragments in mitosis and micronuclei are defective in recruitment of DNA repair proteins.** ***(A)*** Schematic of experimental setup for *(B-D)*. ***(B)*** Immunoblot showing depletion of Rad50 and Mre11 for the experiment described in *(A)*. Quantitation of ***(C)*** dispersed Y-chromosome fragments or ***(D)*** damaged Y chromosomes from the experiments outlined in *(A)* after siRNA depletion of Mre11 or Rad50. (For *(C)* and *(D)*, n=3 independent experiments; total 726, 742, and 727 cells were analyzed for control, siMre11, and siRad50 conditions, respectively. One-way analysis of variance with Bonferroni's multiple comparison test was applied, ns P>0.05.) ***(E)*** Quantitation of damaged Y-chromosomes from the experiment outlined in *(A)* upon siRNA depletion of PolQ (n=3 independent experiments; total 240 and 217 cells were analyzed for control and PolQ depletion, respectively). (Two-tailed unpaired t-test was applied for data shown in *(E)*; ns P>0.05.). Representative images showing lack of recruitment to damaged micronuclei of ***(F)*** 53BP1 or ***(G)*** BRCA1.

**Trivedi et. al. Extended Data Fig. 4. Localization of MDC1, TOPBP1, and CIP2A to micronuclei.**

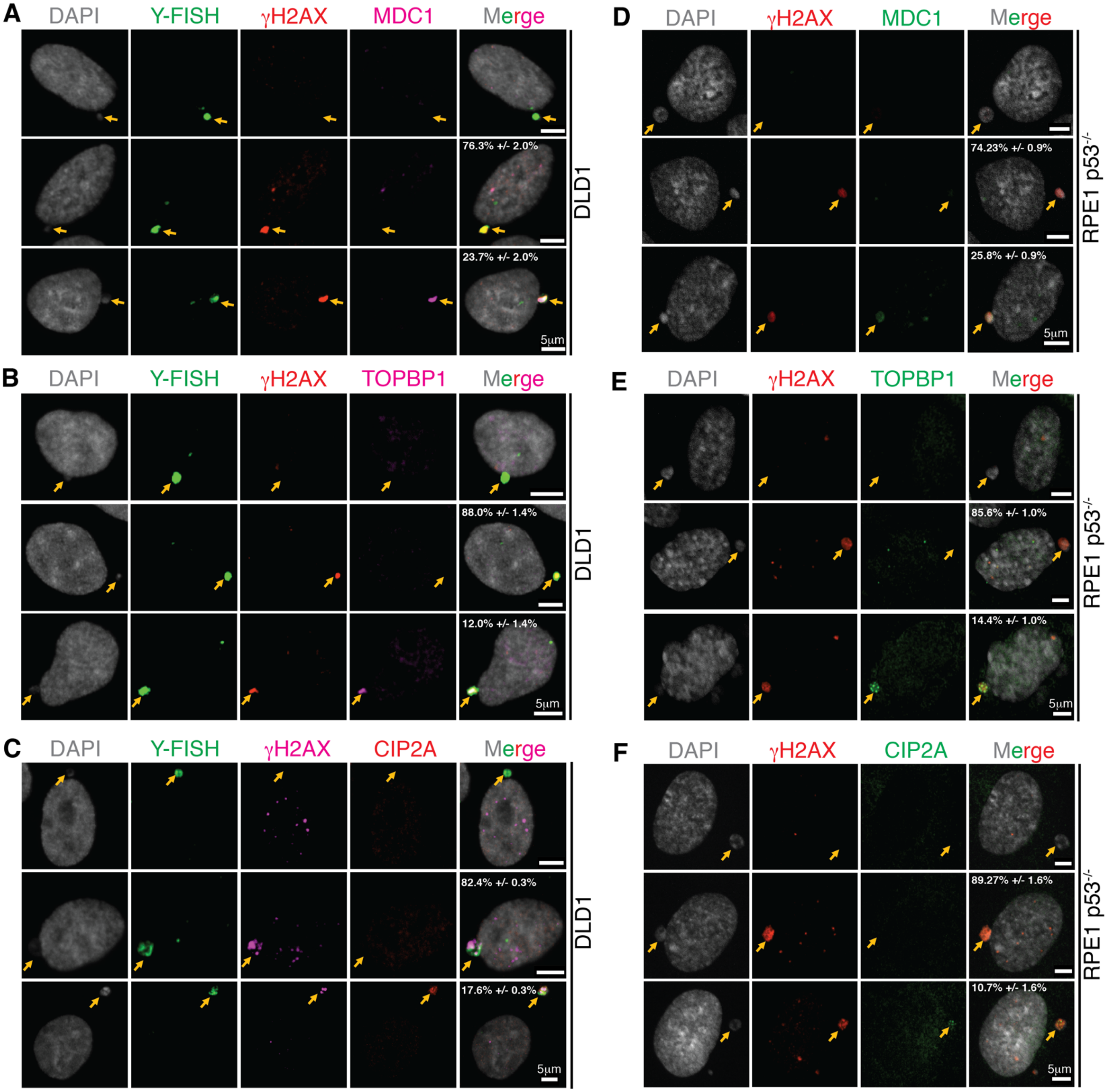


**Extended Data Fig. 4. Localization of MDC1, TOPBP1, and CIP2A to micronuclei.** Representative images showing localization of ***(A, D)*** MDC1, ***(B, E)*** TOPBP1, and ***(C, F)*** CIP2A in micronuclei of DLD1 (n=3 independent experiments, total 482, 290, and 319 DLD1 cells were analyzed for MDC1, TOPBP1, and CIP2A, respectively) and RPE p53$^{-/-}$ (n=3 independent experiments, total 315, 360, and 299 RPE p53$^{-/-}$ cells were analyzed for MDC1, TOPBP1, and CIP2A, respectively) (percentage of cells (mean +/- standard error) with γH2AX positive micronuclei showing the phenotype in the images are indicated on the images). Yellow arrows point to the micronuclei.

**Trivedi et al. Extended Data Fig. 5. Fates of shattered micronuclear chromosomes during mitosis.**

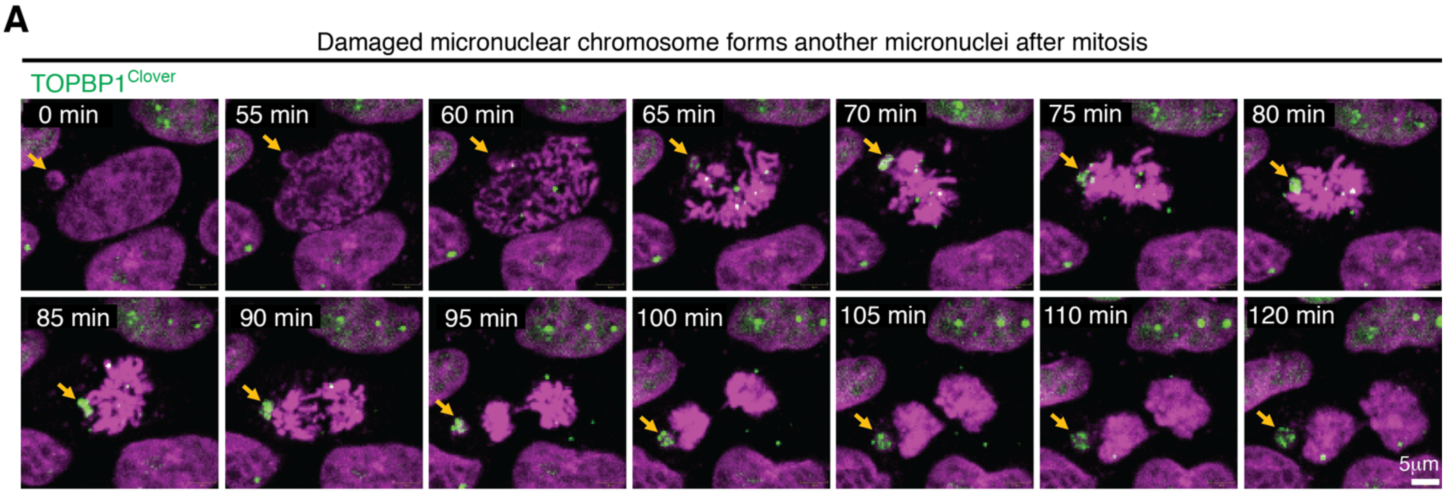


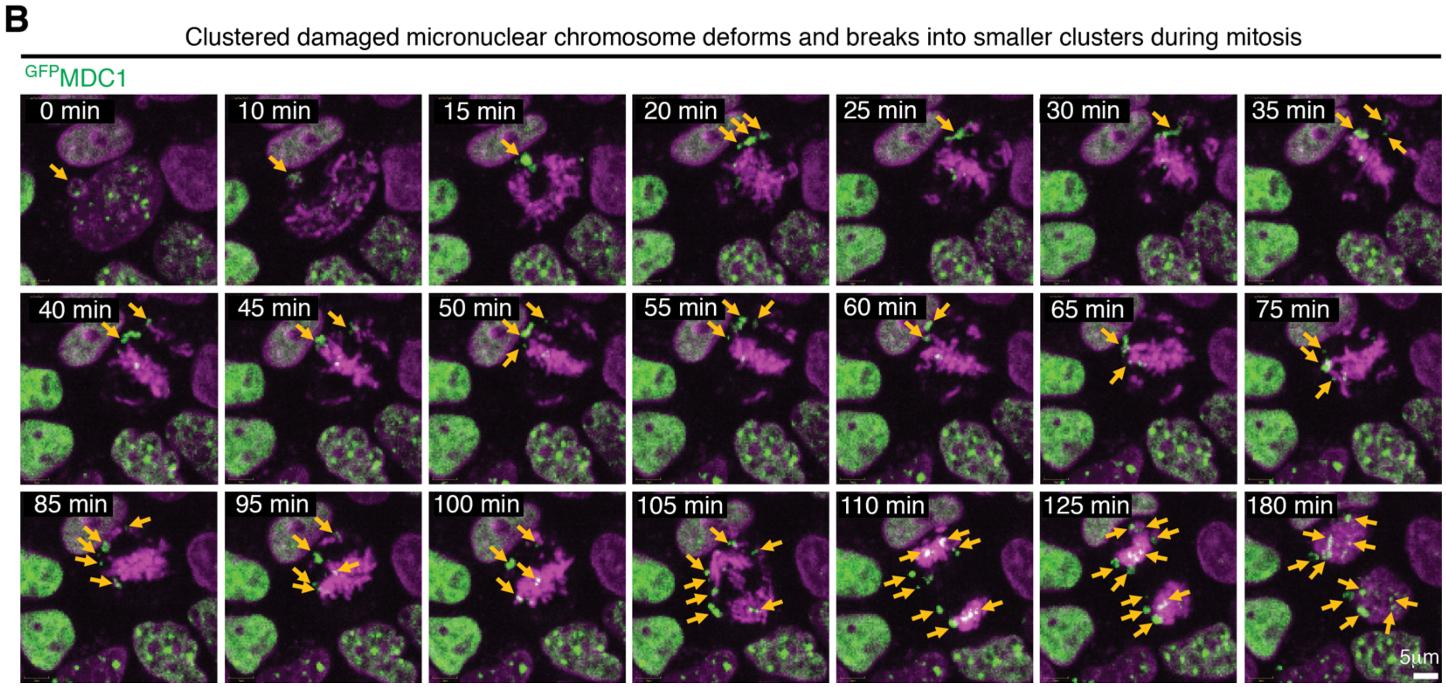


**Extended Data Fig. 5. Fates of shattered micronuclear chromosomes during mitosis.** ***(A and B)*** Frames of **Sup. Videos 3 and 4** showing different behaviors of micronuclear chromosomes throughout mitosis.

**Trivedi et al. Extended Data Fig. 6. Role of MDC1, TOPBP1, and CIP2A in repair of a damaged micronuclear chromosome.**

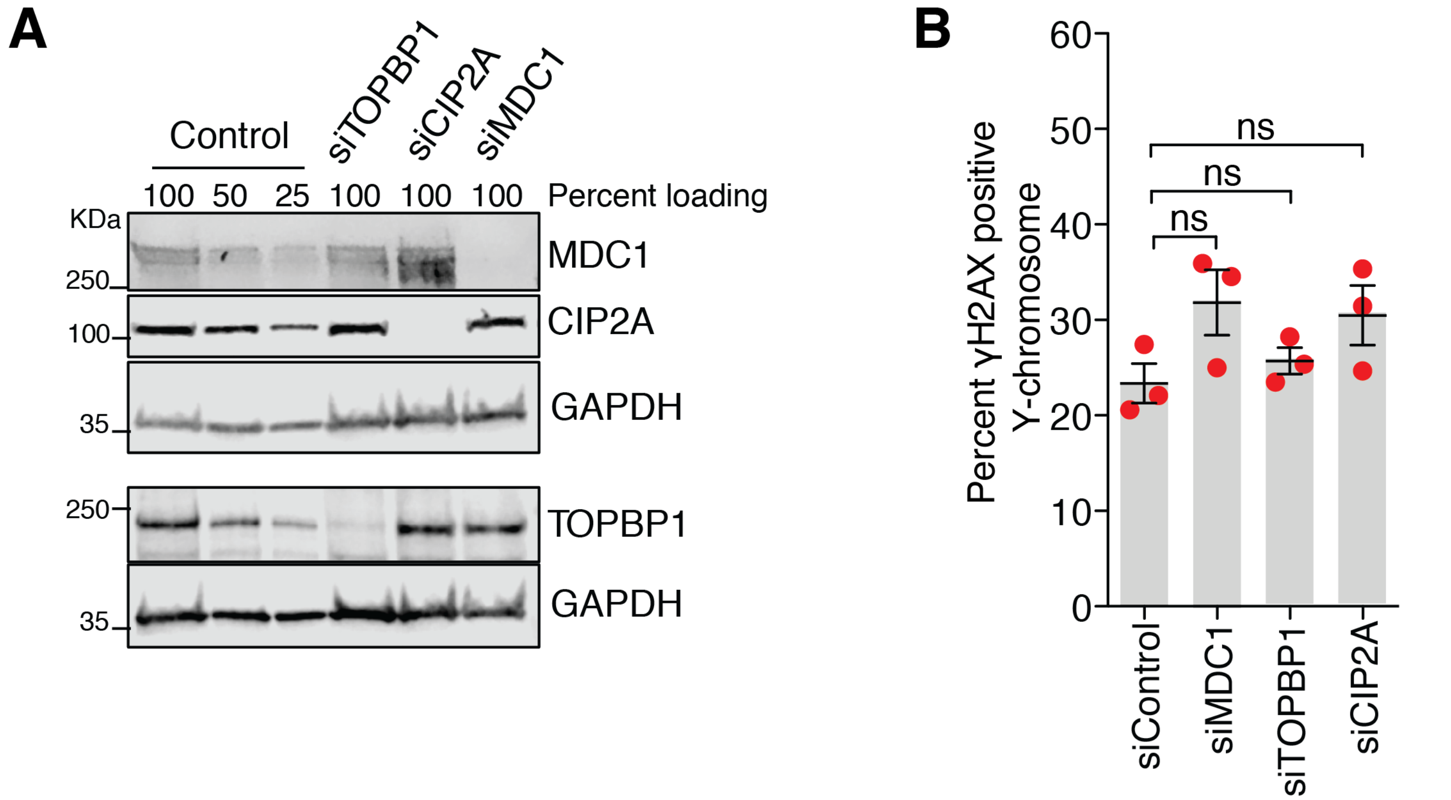


**Extended Data Fig. 6. Role of MDC1, TOPBP1, and CIP2A in repair of a damaged micronuclear chromosome.** ***(A)*** Immunoblot showing depletion of TOPBP1, CIP2A, and MDC1 from the experiment shown in **Fig. 3*A-C***. ***(B)*** Quantitation of damaged Y-chromosomes (tethered or dispersed) for the experiment outlined in **Fig. 3*A*** (n=3 independent experiments; number of cells analyzed is the same as in **Fig. 3*C***). (One-way analysis of variance with Bonferroni's multiple comparison test was applied, ns P>0.05.)

**Trivedi et. al. Extended Data Fig. 7. Role of TOPBP1 in tethering damaged micronuclear chromosomal fragments during mitosis.**

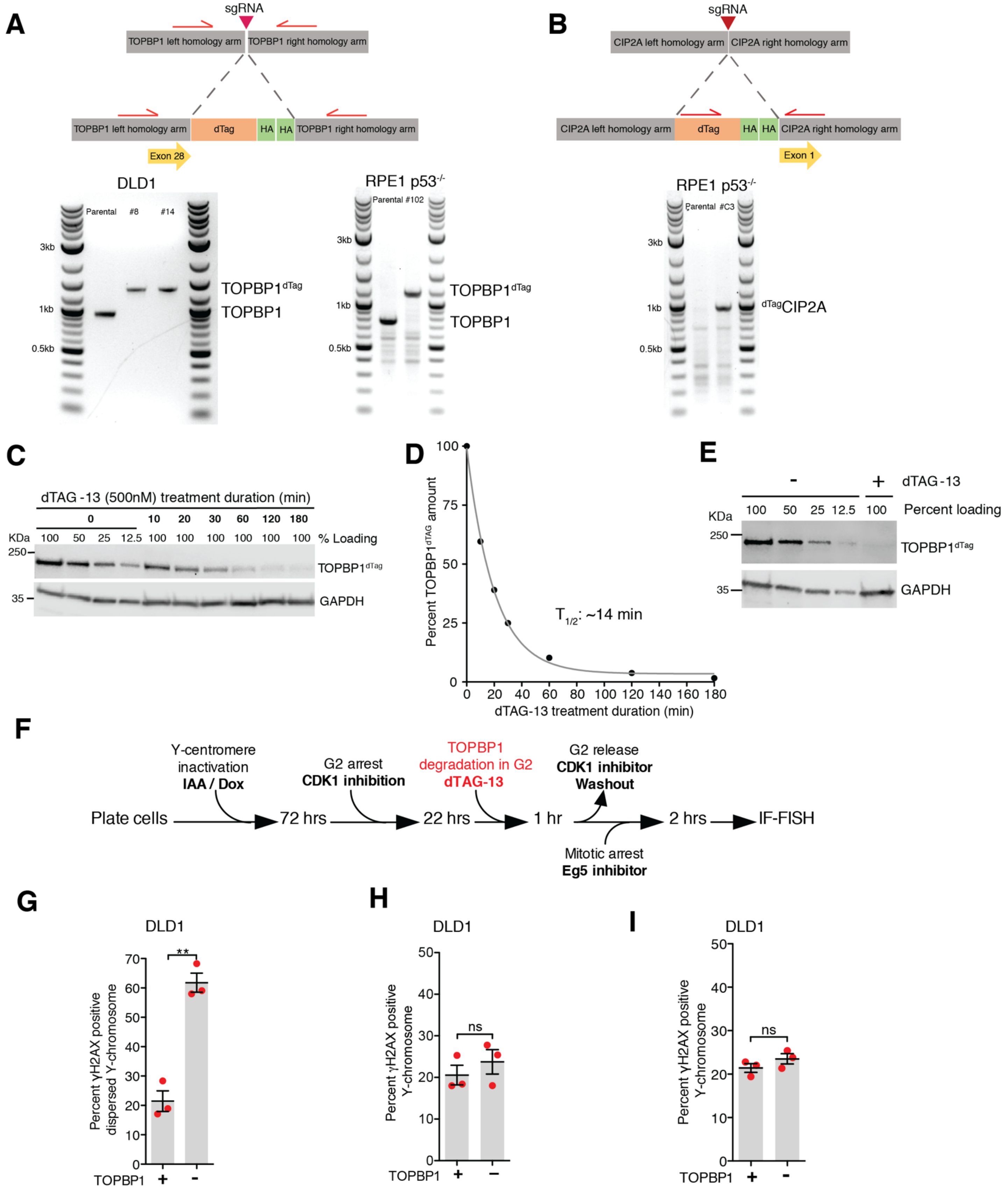


**Extended Data Fig 7. Role of TOPBP1 in tethering damaged micronuclear chromosomal fragments during mitosis.** Schematic of CRISPR-Cas9 mediated tagging of ***(A)*** TOPBP1$^{dTag}$ and ***(B)*** $^{dTag}$CIP2A at both endogenous genomic loci and image of an agarose gel showing successful bi-allelic tagging. ***(C)*** Immunoblot showing dynamics of $TOPBP1^{dTag}$ degradation upon addition of dTAG-13. ***(D)*** Quantitation of $TOPBP1^{dTag}$ levels from (B). Data were fitted with single exponential decay curve, revealing a half-life of 14 minutes. ***(E)*** Immunoblot showing depletion of $TOPBP1^{dTag}$ upon addition of dTAG-13 from the experiment outlined in **Fig. 3*D***. ***(F)*** Experimental outline for *(G)* and *(H)*. ***(G)*** Quantitation of cells with dispersed micronuclear Y-fragments in mitosis upon degradation of $TOPBP1^{dTag}$ in G2 phase. ***(H)*** Quantitation of cells with damaged Y-chromosome fragments (tethered or dispersed) in mitosis after degradation of $TOPBP1^{dTag}$ in G2 phase (for *(G)* and *(H)*, n=3 independent experiments; total 693 and 672 cells were analyzed for control and $TOPBP1^{dTag}$ degradation conditions, respectively). ***(I)*** Quantitation of cells with damaged Y-fragments (tethered or dispersed) in mitosis upon degradation of $TOPBP1^{dTag}$ in mitosis for the experiment outlined in **Fig. 3*D*** (the number of cells analyzed is the same as in **Fig. 3*E***). For *(G-I)*, a two-tailed unpaired t-test was applied; *** P<0.0001 and ns P>0.05).

**Trivedi et al. Extended Data Fig. 8. Role of TOPBP1 and CIP2A in tethering damaged chromosomal fragments resulting from a chromatin bridge during mitosis.**

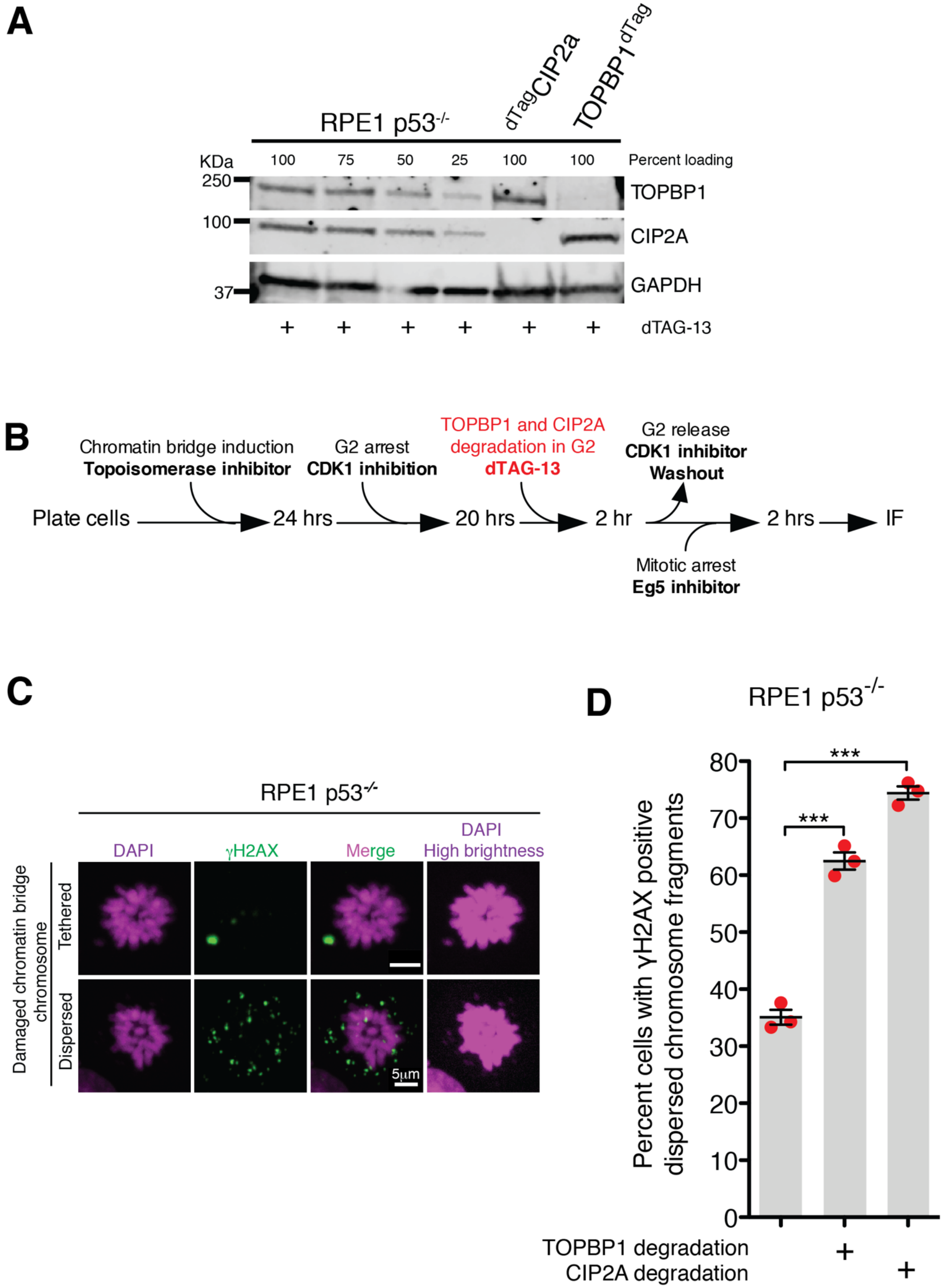


**Extended Data Fig. 8. Role of TOPBP1 and CIP2A in tethering damaged chromosomal fragments resulting from a chromatin bridge during mitosis.** ***(A)*** Immunoblot showing degradation of $^{dTag}$CIP2A and

TOPBP1$^{dTag}$ for the experiment outlined in **Fig. 3*F***. ***(B)*** Experimental outline for *(C)* and *(D)*. ***(C)*** Representative images of RPE p53$^{-/-}$ cells showing tethered and dispersed damaged micronuclear fragments in mitosis after induction of chromatin bridge by inhibition of Topoisomerase II. ***(D)*** Quantitation of cells with dispersed micronuclear fragments in mitosis following chromatin bridge induction upon degradation of TOPBP1$^{dTag}$ or $^{dTag}$CIP2A (n=3 independent experiments; total 420, 518, and 396 cells were analyzed for control, TOPBP1$^{dTag}$ degradation, and $^{dTag}$CIP2A degradation conditions, respectively). (One-way analysis of variance with Bonferroni's multiple comparison test was applied, *** P<0.0001.)

**Trivedi et al. Extended Data Fig. 9. PP2A inhibition by CIP2A is not required for tethering damaged micronuclear chromosomal fragments during mitosis.**

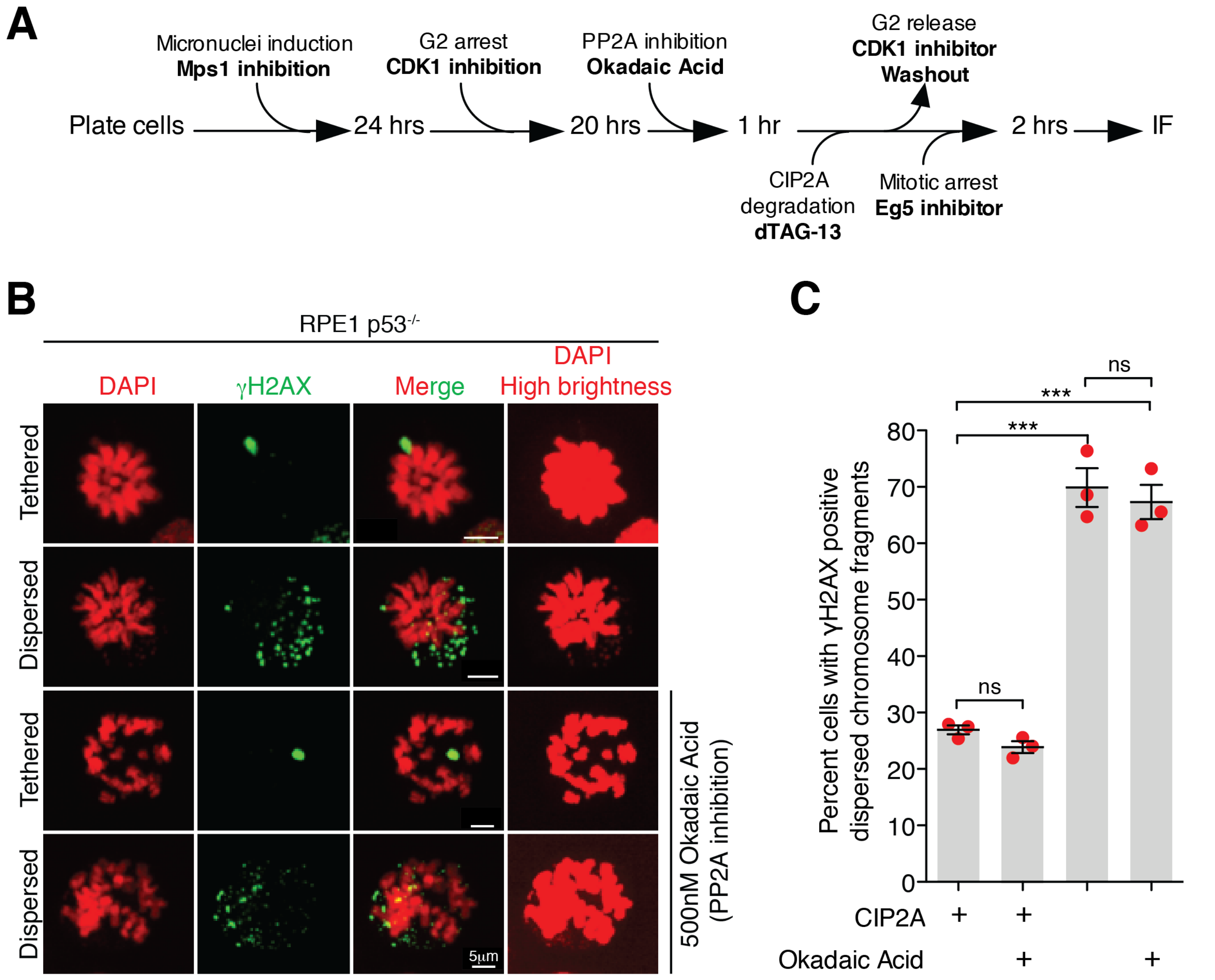


**Extended Data Fig. 9. PP2A inhibition by CIP2A is not required for tethering damaged micronuclear chromosomal fragments during mitosis.** ***(A)*** Experimental outline for the experiment shown in *(B)* and *(C)*. ***(B)*** Representative images of RPE p53$^{-/-}$ cells showing tethered and dispersed damaged micronuclear fragments in mitosis upon the indicated treatment(s). ***(C)*** Quantitation of cells with dispersed micronuclear fragments in mitosis following micronuclei induction upon degradation of $^{dTag}$CIP2A (n=3 independent experiments; total 361, 281, 351, and 364 cells were analyzed for control, okadaic acid treated, $^{dTag}$CIP2A degradation, and $^{dTag}$CIP2A degradation with okadaic acid treated conditions, respectively). (One-way analysis of variance with Bonferroni's multiple comparison test was applied, *** P<0.0001 and ns P>0.05).

**Trivedi et al. Extended Data Fig. 10. High expression of MDC1, TOPBP1, and CIP2A is retained in tumors with a chromothriptically rearranged chromosome.**

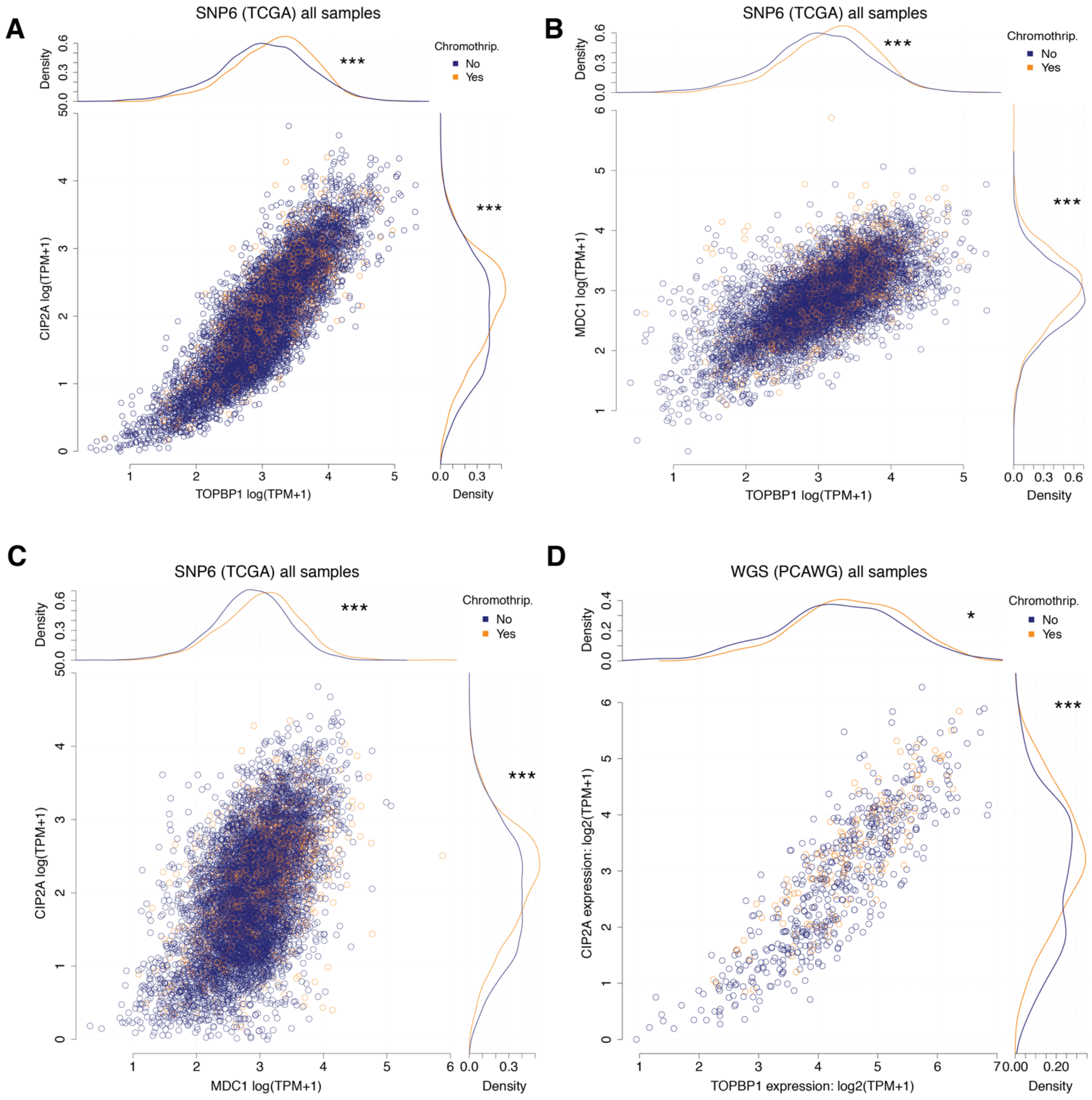


**Extended Data Fig. 10. High expression of MDC1, TOPBP1, and CIP2A is retained in tumors with a chromothriptically rearranged chromosome.** ***(A)*** Associations between gene expression (log(TPM+1)) of TOPBP1 (x-axis) and CIP2A (y-axis) as well as chromothriptic designation of samples (color) for 7,856 tumor samples from The Cancer Genome Atlas (TCGA). Main panel – scatter plot of TOPBP1 and CIP2A expression. Top panel – density of TOPBP1 expression in chromothriptic (orange) and non-chromothriptic samples (blue). Right panel – density of CIP2A expression in chromothriptic and non-chromothriptic samples. Difference in expression between chromothriptic and non-chromothriptic samples is tested with a two-sided Mann-Whitney test; *p<0.05, **p<0.01, ***p<0.001. Chromothripsis is defined by CTLPscanner from SNP-array derived copy number profiles. ***(B)*** Associations between gene expression (log(TPM+1)) of TOPBP1 (x-axis) and MDC1 (y-axis) as well as chromothriptic designation of samples (color) for 7,856 tumor samples from The Cancer Genome Atlas (TCGA). Main panel – scatter plot of TOPBP1 and MDC1 expression. Top panel – density of TOPBP1 expression in chromothriptic (orange) and non-chromothriptic samples (blue). Right panel – density of MDC1 expression in chromothriptic and non-chromothriptic samples. Difference in expression between chromothriptic and non-chromothriptic samples is tested with a two-sided Mann-Whitney test; *p<0.05, **p<0.01, ***p<0.001. Chromothripsis is defined by CTLPscanner from SNP-array derived copy number profiles. ***(C)*** Associations between gene expression (log(TPM+1)) of MDC1 (x-axis) and CIP2A (y-axis) as well as chromothriptic designation of samples (color) for 7,856 tumor samples from The Cancer Genome Atlas (TCGA). Main panel – scatter plot of MDC1 and CIP2A expression. Top panel – density of MDC1 expression in chromothriptic (orange) and non-chromothriptic samples (blue). Right panel – density of CIP2A expression in chromothriptic and non-chromothriptic samples. Difference in expression between chromothriptic and non-chromothriptic samples is tested with a two-sided Mann-Whitney test; *p<0.05, **p<0.01, ***p<0.001. Chromothripsis is defined by CTLPscanner from SNP-array derived copy number profiles. ***(D)*** Associations between gene expression (log(TPM+1)) of TOPBP1 (x-axis) and CIP2A (y-axis) as well as chromothriptic designation of samples (color) for 667 tumor samples from The Cancer Genome Atlas (TCGA) that overlap with Pan-cancer analysis of Whole Genomes (PCAWG) samples. Main panel – scatter plot of TOPBP1 and CIP2A expression. Top panel – density of TOPBP1 expression in chromothriptic (orange) and non-chromothriptic samples (blue). Right panel – density of CIP2A expression in chromothriptic and non-chromothriptic samples. Difference in expression between chromothriptic and non-chromothriptic samples is tested with a two-sided Mann-Whitney test; *p<0.05, **p<0.01, ***p<0.001. Chromothripsis is defined by shatterSeek from WGS-derived copy number profiles.

**Trivedi et al. Extended Data Fig.11. Positive associations of Cip2A and TOPBP1 expression with extent of loss of heterozygosity (LOH).**

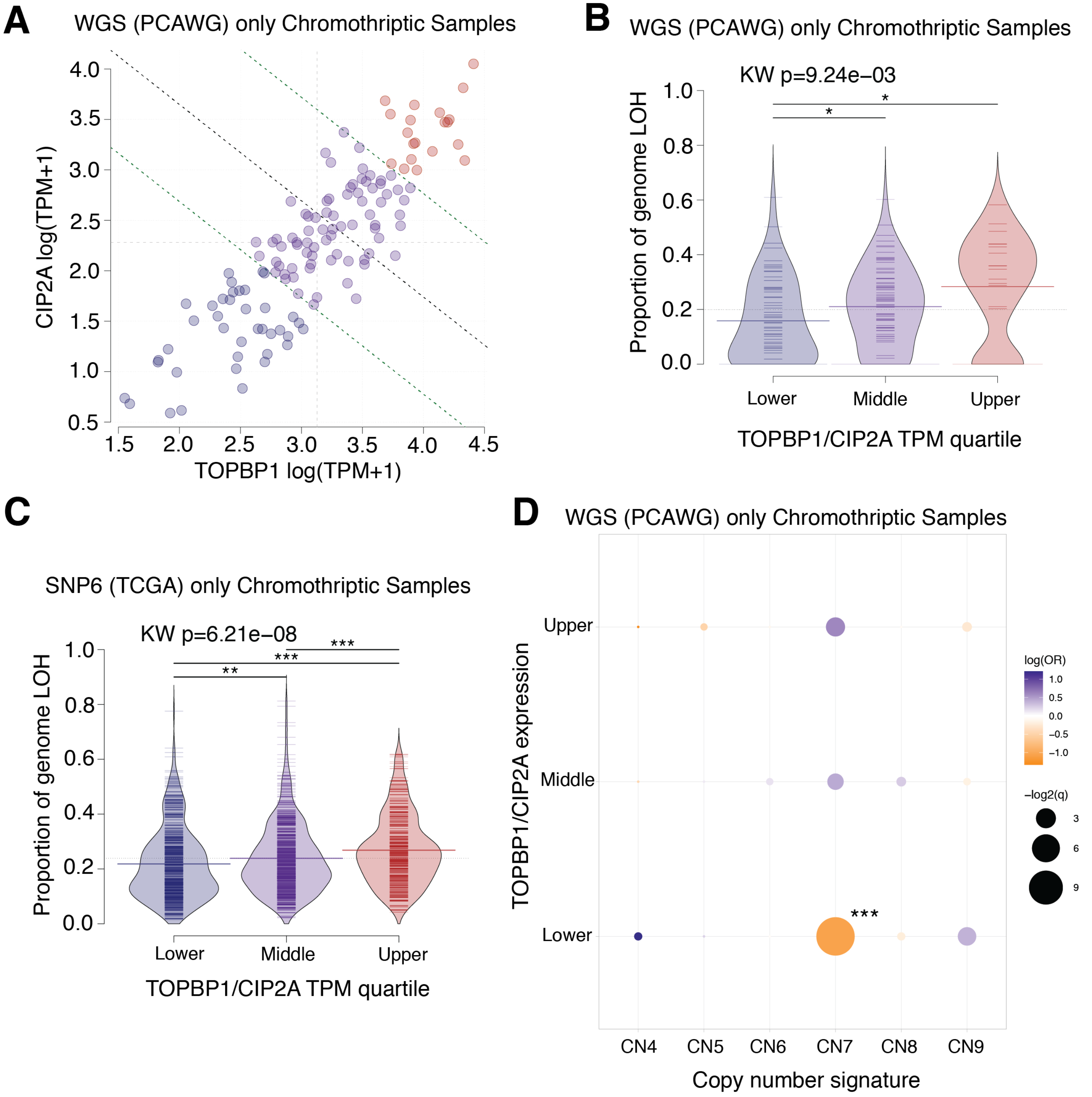


**Extended Data Fig.11. Positive associations of Cip2A and TOPBP1 expression with extent of loss of heterozygosity (LOH).** ***(A)*** Gene expression of TOPBP1 and CIP2A (log(TPM+1)) in 135 chromothriptic tumor samples from The Cancer Genome Atlas (TCGA) that overlap with Pan-cancer Analysis of Whole Genomes (PCAWG) samples. Samples are categorized as joint upper (red), middle (purple) or lower (blue) expression of both genes by bisecting the line of best fit with a decision boundary at the upper quartile or lower quartiles of each gene (green dotted lines). Light grey dotted lines denote median values of gene expression, black dotted line indicates the bisecting line of the median. TPM=transcripts per million. Chromothriptic samples were determined by shatterSeek. ***(B)*** Association between the proportion of genome designated as loss of heterozygosity (LOH; y-axis) and gene expression groups defined in *(A)*. Thick colored horizontal lines indicate mean proportion of LOH for each expression group. Thin horizontal lines indicate individual samples. Overall significance was tested with a Kruskal-Wallis test. Pairwise comparisons were performed with two-sided Mann-Whitney tests. *p<0.05. KW=Kruskal-Wallis test. ***(C)*** Association between the proportion of genome designated as loss of heterozygosity (LOH; y-axis) and gene expression groups defined in **Fig. 4*H*** for chromothriptic samples only. Thick coloured horizontal lines indicate mean proportion of LOH for each expression group. Thin horizontal lines indicate individual samples. Black horizontal line indicates the mean proportion of LOH across the dataset. Overall significance was tested with a Kruskal-Wallis test. Pairwise comparisons were performed with two-sided Mann-Whitney tests. **p<0.01, ***p<0.001. KW=Kruskal-Wallis test. Chromothripsis is defined by CTLPscanner from SNP-array derived copy number profiles. ***(D)*** Associations between gene expression groups defined in *(A)* (y-axis), and chromothripsis associated copy number signatures (x-axis; CN4:9), using two-sided Fisher's exact tests. Correction for multiple testing was performed using the Benjamini-Hochberg method. Color denotes odds ratio, size of points indicates the significance of the test. *q<0.05, **q<0.01, ***q<0.001. OR=odds ratio. CN7 is a signature associated with chromothripsis amplification.

**Trivedi et al. Extended Data Fig. 12. Model for chromothriptic chromosome formation.**

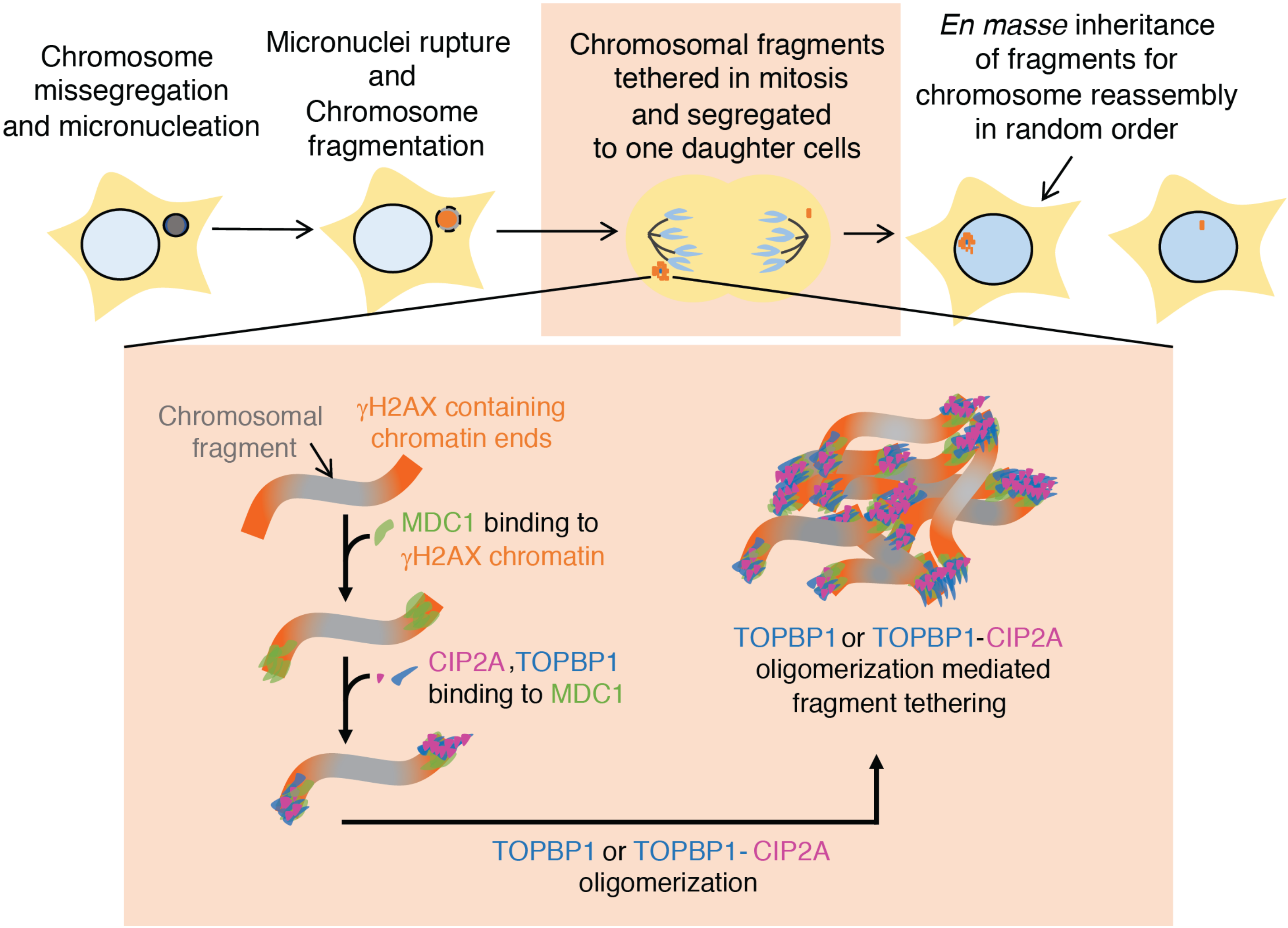


**Extended Data Fig. 12. Model for chromothriptic chromosome formation.** Chromosomes within micronuclei (that were formed as a result of errors in mitosis) are fragmented before or upon entry into mitosis. The fragments are tethered during mitosis by combined action of MDC1, TOPBP1, and CIP2A. Tethering of chromosomal fragments during mitosis ensures their transfer to the same daughter nucleus for subsequent ligation in random order, thereby resulting in reassembly of a heritable, highly rearranged chromosome.

**Supplement Video 1.** DLD1 cell with micronucleus, induced by Y-centromere inactivation, expressing $^{GFP}$MDC1 going through mitosis.

**Supplement Video 2.** DLD1 cell with micronucleus, induced by Y-centromere inactivation, expressing TOPBP1$^{Clover}$ going through mitosis.

**Supplement Video 3.** DLD1 cell with micronucleus, induced by Y-centromere inactivation, expressing TOPBP1$^{Clover}$ going through mitosis. The damaged micronuclear chromosomal fragments remains tethered and form a new micronucleus in the daughter cell.

**Supplement Video 4.** DLD1 cell with micronucleus, induced by Y-centromere inactivation, expressing $^{GFP}$MDC1 going through mitosis. The damaged micronuclear chromosomal fragments form a cluster initially which deforms and breaks into smaller cluster during passage through mitosis.

**Supplement Video 5.** RPE p53$^{-/-}$ TOPBP1$^{dTag}$ cell, arrested in mitosis by Eg5 inhibitor, with damaged and clustered chromosome induced by Mps1 inhibition and marked with $^{GFP}$MDC1. TOPBP1 degradation is not induced and the chromosomal fragments remain clustered during mitosis.

**Supplement Video 6.** RPE p53$^{-/-}$ TOPBP1$^{dTag}$ cell, arrested in mitosis by Eg5 inhibitor, with damaged and clustered chromosome induced by Mps1 inhibition and marked with $^{GFP}$MDC1. TOPBP1 degradation was induced (by addition of dTAG-13) at the start of the video, the chromosomal fragments are clustered initially but disperse as TOPBP1$^{dTag}$ is degraded.
